# Semi-rational evolution of a recombinant DNA polymerase for modified nucleotide incorporation efficiency

**DOI:** 10.1101/2023.03.20.533374

**Authors:** Lili Zhai, Zi Wang, Fen Liu, Chongjun Xu, Jingjing Wang, Hongyan Han, Qingqing Xie, Wenwei Zhang, Yue Zheng, Alexander K. Buell, Yuliang Dong

## Abstract

Engineering improved B-family DNA polymerases to incorporate 3′-O-modified nucleotide reversible terminators is limited by an insufficient understanding of the structural determinants that define polymerization efficiency. To explore the key mechanism for unnatural nucleotide incorporation, we engineered a B-family DNA polymerase from *Thermococcus Kodakaraenis* (KOD pol) by using semi-rational design strategies. We first scanned the active pocket of KOD pol through site-directed saturation mutagenesis and combinatorial mutations and identified a variant Mut_C2 containing five mutation sites (D141A, E143A, L408I, Y409A, A485E) using a high-throughput microwell-based screening method. Mut_C2 demonstrated high catalytic efficiency in incorporating 3’-O-azidomethyl-dATP labeled with a Cy3 dye, whereas the wild-type KOD pol failed to incorporate it. Computational simulations were then conducted towards the DNA binding region of KOD pol to predict additional mutations with enhanced catalytic activity, which were subsequently experimentally verified. By a stepwise combinatorial mutagenesis approach, we obtained an eleven-mutation variant, named Mut_E10 by introducing additional mutations to the Mut_C2 variant. Mut_E10, which carried six specific mutations (S383T, Y384F, V389I, V589H, T676K, and V680M) within the DNA-binding region, demonstrated over 20-fold improvement in kinetic efficiency as compared to Mut_C2. In addition, Mut_E10 demonstrated satisfactory performance in two different sequencing platforms (BGISEQ-500 and MGISEQ-2000), indicating its potential for commercialization. Our study demonstrates that an effective enhancement in its catalytic efficiency towards modified nucleotides can be achieved efficiently through combinatorial mutagenesis of residues in the active site and DNA binding region of DNA polymerase. These findings contribute to a comprehensive understanding of the mechanisms that underlie the incorporation of modified nucleotides by DNA polymerase. The beneficial mutation sites, as well as the nucleotide incorporation mechanism identified in this study, can provide valuable guidance for the engineering of other B-family DNA polymerases.

## Introduction

DNA polymerases discovered in nature play important roles in molecular biology, synthetic biology, and molecular diagnostics^1,2^. Wilde-type (WT) DNA polymerases mostly fail or perform poorly in practical applications^2^. However, these shortcomings can be overcome by enzyme engineering^3^. The application of DNA polymerase engineering to improve the performance of natural enzymes has been widely reported^1,4^. Engineered DNA polymerases are the workhorses of numerous biotechnological applications such as in the fields of DNA synthesis^4^, molecular diagnostics^5^, and DNA sequencing^3,6,7^.

Many DNA sequencing methods rely on engineered DNA polymerases that efficiently incorporate chemically modified nucleotides^8,9^ such as the Sanger sequencing method, which relies on dideoxynucleoside triphosphates (ddNTPs)^7^, and the sequencing-by-synthesis (SBS) method, which uses dye-labeled reversible terminators in an iterative loop^6^. SBS relies on the identification of each base as the DNA strand is extended by the cleavable fluorescent nucleotide reversible terminators that temporarily pause the DNA synthesis for sequence determination^10^. Challenges in using SBS with cleavable fluorescent nucleotide reversible terminators involve further improving the DNA polymerase that efficiently recognizes and incorporates the modified nucleotides^9^. Archaeal B-family DNA polymerases are widely engineered to be applied in catalyzing the addition of chemically modified nucleotide substrates in SBS sequencing, because of their potential for tunability of the active site and acceptance of broad substrate^11,12,13^. The engineering of B-family DNA polymerases relies on the development of various enzyme evolution techniques^14^.

Enzyme evolution techniques usually include computational design^15,16^, semi-rational design^17^, and large-scale screening by directed evolution^18^. Semi-rational design is one of the most widely applied approaches, which enables targeted exploration of functional amino acid sites by selecting and mutating specific residues, with reduced library size and screening costs compared to random mutagenesis^14^. Semi-rational design is typically complemented by various screening methods to evaluate and select protein variants with desired characteristics^17^. Enzyme screening strategies include DNA shuffling^19^, CSR (Compartmentalized Self-Replication)^20^, microfluidic screening technology^21^, microplate reader screening technology^22^, etc. Most screening techniques will utilize fluorescence as a readout because of its high sensitivity, time efficiency, the potential for high throughput, low cost, and other advantages^23^. Förster resonance energy transfer (FRET) is a mechanism describing energy transfer between two fluorescent molecules^24^, which is widely exploited in various protein mechanistic and engineering studies, such as studies of protein folding kinetics^25^, exploring the spatial relationships between protein and substrates^26^, and protein-protein interactions^26^. In particular, previous studies have reported the applicability of FRET technology for the screening of enzymatic activity^27,28^.

KOD DNA polymerase (KOD pol) originates from *Thermococcus kodakarensis* which is an extremophilic archaeon, an important representative of B-family DNA polymerases^29,30^. Engineered KOD pol is the key component of numerous biotechnological applications such as multiple types of PCR^31^, TNA synthesis^32^, and incorporation of base or sugar-modified nucleoside triphosphates^33,34,29^. Some KOD variants have been reported to tolerate and polymerize certain unnatural nucleotides^35^. For example, the KOD variant (exo-) can tolerate 7-deaza-modified adenosine triphosphate^36^ or microenvironment-sensitive fluorescent nucleotide probes^34^. KOD variant (exo-, A485L) can incorporate 5-substituted pyrimidine nucleoside triphosphates (dNamTPs)^37^. KOD variant (P179S, L650R) can tolerate LNA nucleotides additionally modified at position 3’ of the sugar moiety^38^. However, significantly less work has been reported on the engineering and modification of B family DNA polymerases to enable the efficient incorporation of cleavable fluorescent nucleotide reversible terminators in Next-Generation Sequencing (NGS) applications. A previous study reported that the 9°N DNA polymerase variant (exo-, A485L, Y409V) can incorporate nucleotide reversible terminators, such as 3’-O-azidomethyl-dNTPs^9^. KOD DNA polymerase and 9°N DNA polymerase share high sequence similarity (91%), and both belong to the B-family polymerases^39^. Thus, KOD polymerase has the potential to be engineered to efficiently incorporate reversible terminators, thereby expanding its applications in the field of DNA sequencing^39^. In Engineering KOD DNA polymerases to incorporate modified nucleotides, there are generally two main directions of sequence modification: mutations in the active site to enable discrimination and incorporation of specific nucleotides^40^, and mutations in the DNA strand binding region to enhance catalytic efficiency by stabilizing the interactions between the polymerase, the DNA template, and the incoming nucleotide^41^.

In this study, we established a FRET-based enzyme screening platform to screen mutants for their improved abilities to incorporate 3’-O-azidomethyl-dATP labeled with a Cy3 dye into a primer-templated and Cy5-labeled DNA using the FRET emission signal of Cy5 as an indicator. Initially, we scanned amino acids located in the active site pocket of KOD pol. Subsequently, we computationally predicted relevant residues to be replaced within the DNA binding region of KOD pol. By employing enzyme kinetic screening of multiple mutant libraries, we successfully obtained a KOD variant that exhibited satisfactory performance on both BGI and MGI sequencing platforms. Through dissecting the specific residues within the active site and DNA binding region, this study provides insights into the key mechanisms underlying the incorporation of modified nucleotides by KOD pol. The effectiveness of semi-rational design strategies and the identified beneficial mutation sites in this study serve as valuable guidance for the engineering of other B-family DNA polymerases. In addition, the significance of our research lies in its implications for the engineering of DNA polymerases and their applications in NGS.

## Results

### Mutant screening strategy establishment

The comprehensive screening strategy developed for KOD pol involved multiple intricate steps including structural analysis, MD simulations, protein expression, mutant screening, mutant protein purification, and tests in sequencing applications, as illustrated in Figure 1A. The process began with a structural analysis of KOD pol to provide valuable insights into potential sites for mutagenesis. We performed saturation and combinatorial mutagenesis on the selected sites to construct multiple libraries. We employed MD simulations to predict more promising sites to do stepwise combinatorial mutagenesis. The high-throughput expression of KOD pol variants was carried out in a 96-deep-well plate format in order to obtain sufficient quantities of protein for screening. The expressed proteins were subsequently subjected to semi-purification followed by quantification to ensure the reliability and comparability of the subsequent mutant screening experiments (Supplementary Figure 1A and Supplementary Figure 1B). Then, the protein variants were screened for enzyme activity and kinetic performance in a 384-well plate using a microplate reader. After identifying variants with favorable kinetic characteristics, we performed rigorous purification using chromatography with three prepacked columns to eliminate potential interference from impurities on sequencing quality. Subsequently, the promising and highly purified proteins were tested for their sequencing performance on the BGISEQ-500 platform for SE50 testing and the MGISEQ-2000 platform for PE100 testing.

**Figure 1.**
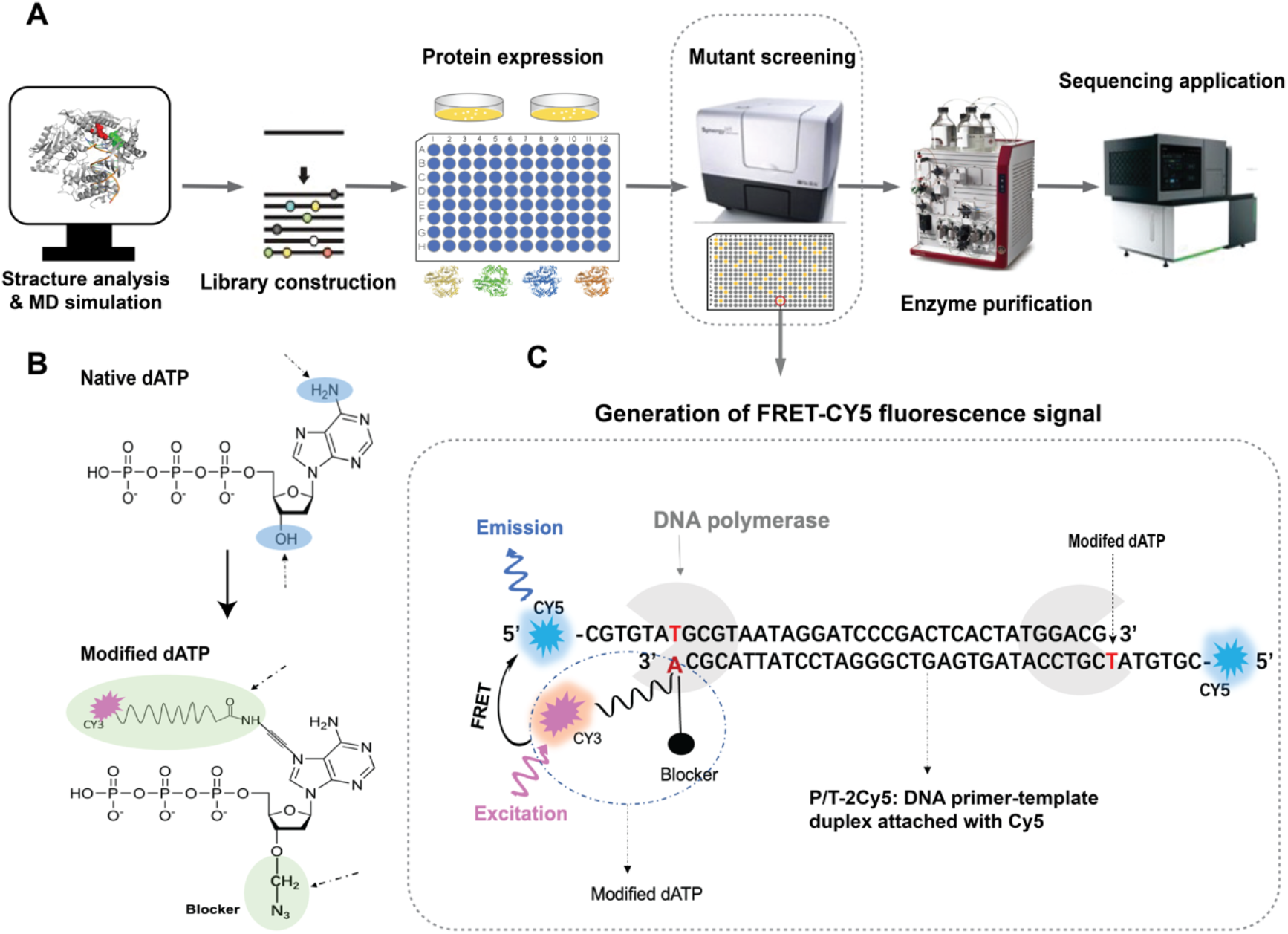
Schematic of KOD pol engineering experiments. A) Overview of the semi-rational evolution process involving structural analysis of KOD pol, MD simulation, protein expression, variant screening, variant purification, and sequencing application tests. Multiple libraries were constructed by site-direct mutagenesis, combinatorial mutagenesis, and stepwise combinatorial mutagenesis. The screening of the catalytic efficiency of the mutants was performed by monitoring FRET Cy5 fluorescence as an indicator. B) The molecular structures of 3’-O-azidomethyl-deoxyadenosine triphosphate with dye Cy3 labeled (modified dATP) as well as native dATP. C) The process of the generation of the FRET Cy5 fluorescence emission signal. The FRET Cy5 fluorescence emission signal can be detected at 676 nm upon excitation of the incorporated Cy3 dye-labeled modified dATP at 530 nm. If the modified dATP is not properly incorporated into the P/T-2Cy5, no FRET Cy5 fluorescence emission signal is detected. DNA polymerase is shown in cartoon format and colored grey. Dye Cy5 and Cy3 are shown in cartoon format and colored blue and pink, respectively. The primer-template duplex is labeled with Cy5 dye at both 5’ ends, which enhances signal intensity.

In the screening method, the modified dATP, 3’-O-azidomethyl-dATP labeled with a Cy3 dye, shown in Figure 1B, was employed. Additionally, we employed a primer-template complex in which the single oligo strand was labeled with a fluorescent Cy5 dye at its 5’ end (P/T-2Cy5), as illustrated in Figure 1C. Upon successful incorporation of 3’-O-azidomethyl-dATP labeled with a Cy3 dye into P/T-2Cy5 by DNA polymerase, excitation at 530 nm of Cy3 results in fluorescence resonance energy transfer to the nearby Cy5 molecule, resulting in the emission at 676 nm by Cy5, as illustrated in Figure 1C. This increase in Cy5 FRET emission signal can be conveniently monitored inside a microplate reader, allowing for the measurement of the rate of modified nucleotide incorporation into the DNA strand. By comparing the incorporation rates (RFUs/min, with RFUs representing the increased Cy5 FRET emission signal), we were able to determine the catalytic efficiency of KOD variants for incorporating modified dATP.

Experimental validation for the screening method was performed successfully by comparing two KOD variants with distinguishing enzymatic activities, and WT KOD pol, shown in Supplementary Figure 1C. The developed screening method was effective in distinguishing between the different levels of enzymatic activities of the variants.

### Selection of first-round mutation sites and mutant library construction

First, we analyzed the amino acids located in the active pocket of KOD pol. The crystal structures of KOD pol, including the open (PDB ID: 4K8Z)^42^ and the closed (PDB ID:5OMF)^39^ states, are superimposed and shown in Figure 2 A. A significant movement of the finger domain from the open state to the closed state can be observed, resulting in the formation of the active pocket by the palm and closed finger subdomains. Residues located in the active pocket of KOD pol directly impact nucleotide incorporation efficiency by interfacing with incoming nucleotides. In the active pocket, residues L408, Y409, P410, and A485 (Figure 2B) were reported to associate with the enhanced unnatural nucleotide incorporation efficiency^43^. In addition, in the engineering of B-family polymerases to incorporate modified nucleotides, exonuclease activity is often inactivated to prevent the excision of the incorporated modified nucleotides^44^. A previous study reported that an exonuclease-deficient variant (D141A and E143A) with A485L mutation derived from *Thermococcus kodakaraensis* (KOD) DNA polymerase was prepared to incorporate unnatural nucleotides into a primer-template DNA^45^. Based on these findings, we selected residues L408, Y409, P410, and A485, as well as mutation sites D141A and E143A with exonuclease deficiency, as key sites for the first-round mutagenesis. Despite the fact that these positions have been previously reported to be important for the objectives of our study, there is still a lack of in-depth investigation into the effects of their combination and interactions with other related sites. Such an in-depth investigation will be valuable for the engineering of B-family DNA polymerases to improve the incorporation efficiency for unnatural nucleotides.

**Figure 2.**
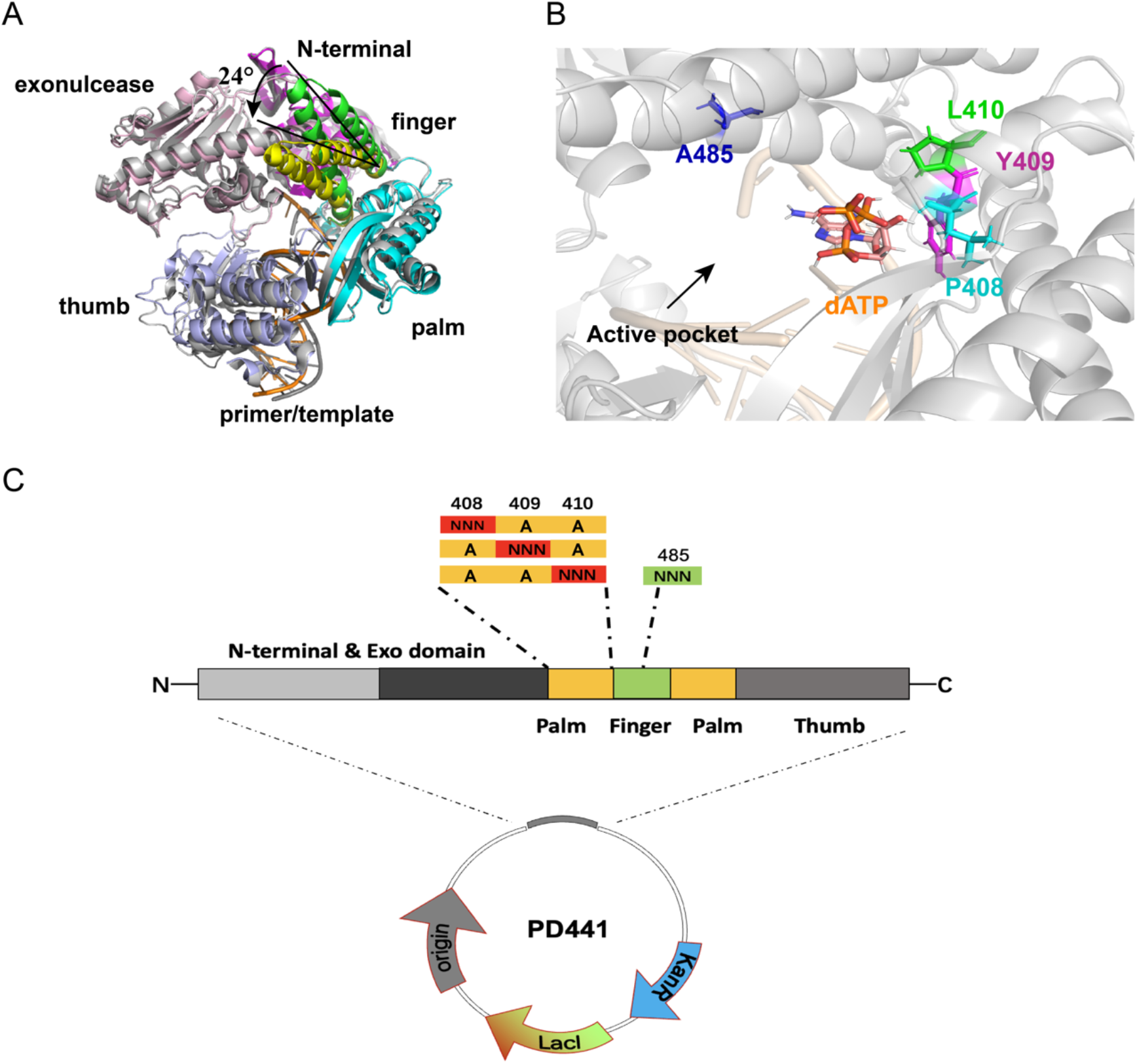
Illustration of structures of KOD pol and demonstration of library construction. A) Depiction of the superimposed structures of a ternary KOD complex (closed state, PDB ID: 5MOF) and a binary KOD complex (open state, PDB ID: 4K8Z), which are shown in cartoon format. The finger domain of the ternary complex is shown in yellow and closed by an inward movement of approximately 24°, in comparison to the finger domain of the binary complex shown in green. The other domains of the KOD ternary complex are colored in gray, the exonuclease of the binary complex in light pink, the N-terminal of the binary complex in pink, the thumb domain of the binary complex in light blue, the palm domain of the binary complex in cyan, and the primer-template of the KOD ternary and binary complex are colored in gray and orange, respectively. B) Illustration of the active pocket of KOD pol with position A485 colored in blue, L408 colored in cyan, Y409 colored in pink, and P410 colored in green. All residues are shown as sticks. The dATP is shown as sticks and depicted in orange elements, and the rest of the protein is shown in cartoon format and colored in gray. C) Illustration of the first-round constructed library, which includes saturation mutations at positions 408, 409, 410, and 485. The plasmid is pD441 which is a high copy number vector for the expression of *E. coli* flagellin from an IPTG-inducible T5 promoter.

We performed alanine substitutions at residues L408, Y409, P410, and A485. Introducing alanine substitutions at critical positions in proteins represents a potent strategy for identifying the residues that play crucial roles in protein function, stability, and structure^46^. Alanine substitution allows for the evaluation of the individual contributions of each amino acid sidechain to the overall functionality of the protein^46,47^. In this study, alanine substitution in the active pocket of KOD pol could create favorable conditions for the incorporation of modified dATP carrying sterically demanding groups, such as a flexible linker and fluorescence dye. Taking inspiration from this approach, we designed a variant of KOD DNA polymerase called Mut_1 (D141A, E143A, L408A, Y409A, P410A), as the parent variant in this round of mutagenesis. Notably, position 485 in the wild-type KOD pol is already an A, so no further mutation was introduced at this site in Mut_1. Next, we constructed the first-round library based on Mut_1, which involved saturation mutagenesis at positions 408, 409, 410, and 485. The mutant library was constructed (Figure 2C) by performing saturation mutagenesis at position 408 with Y409A/P410A, saturation mutagenesis at position 409 with L408A/P410A, saturation mutagenesis at position 410 with L408A/Y409A, and saturation mutagenesis at position 485 in the case of the mutant with L408A/Y409A/P410A. These libraries were subjected to enzyme activity screening to investigate the impact of specific residue positions on the catalytic efficiency, which laid the groundwork for subsequent combinatorial mutagenesis.

### Mutant screening of the first-round mutagenesis and combinatorial (the second-round) mutagenesis

The screening results of the first-round library are presented in Figure 3. The data were shown as relative reaction velocity, V_rel_ as a measure for the enzyme activity of KOD mutants. V_rel_ of KOD variants was calculated with equation 1:

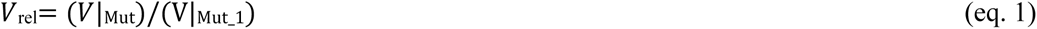

**Figure 3.**
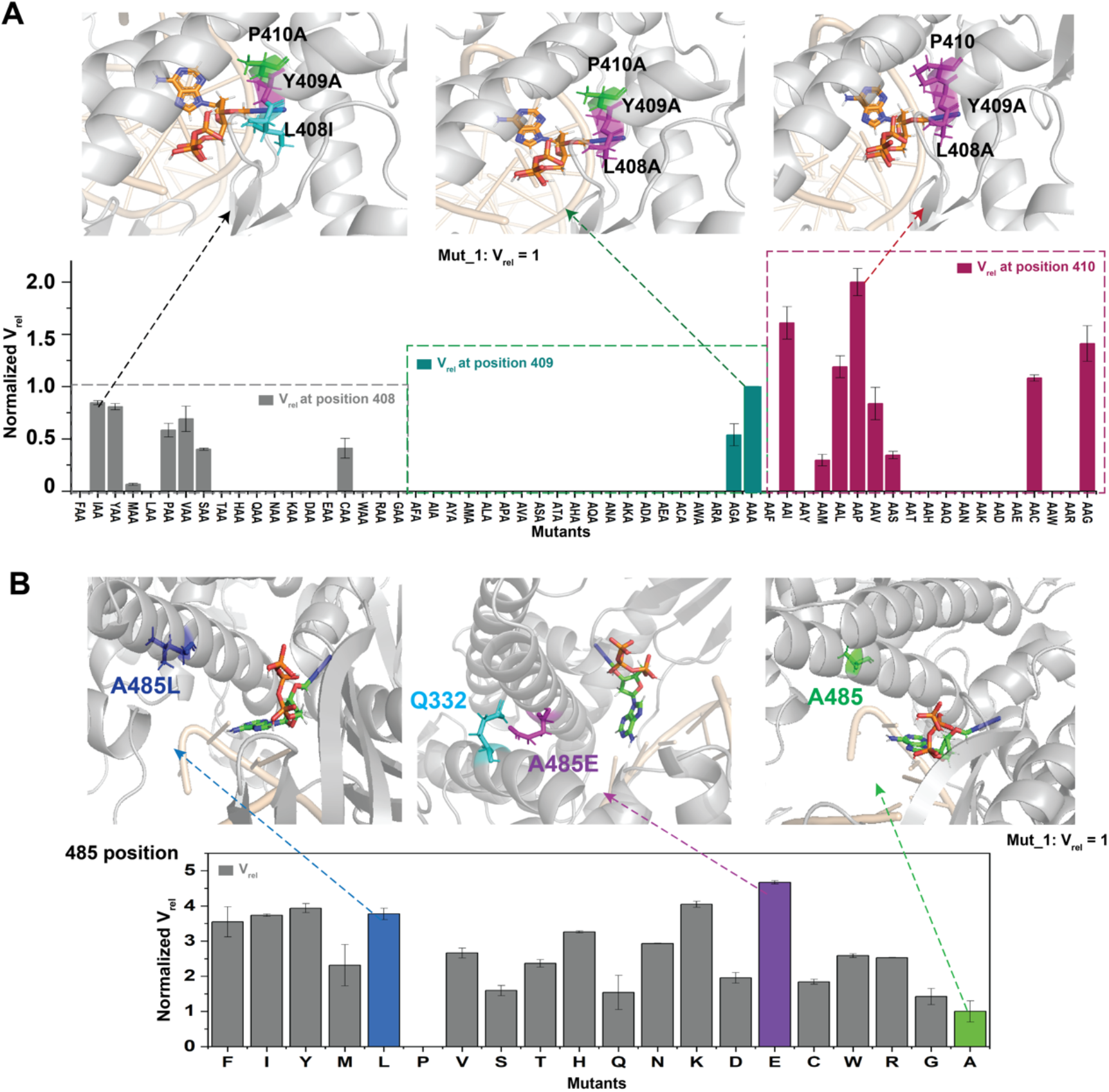
Summary of the screening results of saturation mutagenesis at positions 408, 409, 410, and 485 of KOD pol. A) The screening results of amino acid substitutions at positions 408, 409, and 410, are depicted as V_rel_ values. The bar charts show the screening results of saturation mutagenesis in positions 408 (gray), 409 (dark green), and 410 (red). The upper panels show locations of L408I (cyan), Y409A (pink), and P410A (green) amino acids as well as 3’-O-azidomethyl-dATP (orange) in the crystal structure of KOD pol (PDB ID: 5MOF). The original dATP was replaced with 3’-O-azidomethyl-dATP, which was positioned using homology modeling. Subsequent mutagenesis experiments were performed using PyMOL. L408I (pink), Y409A (pink), P410A (green), and 3’-O-azidomethyl-dATP (orange) are shown as sticks. The rest of the protein structure is shown in cartoon format and colored grey, and the DNA strand is shown as cartoon and colored in wheat. B) The screening results of saturation mutagenesis at position 485, as V_rel_ like in A). The blue bar represents the A485L substitution, with the corresponding structure shown above. The purple bar represents the A485E substitution, with the corresponding structure shown above and an observed interaction with Q332 (cyan). The green bar represents the original A485 residue and its corresponding structure.

V|_Mut_1_ represents the incorporation rate V (RFUs/min) of Mut_1, while *V|*_Mut_ represents the incorporation rate V (RFUs/min) of the given variant. The parent variant Mut_1 had a V_rel_ value of 1, as shown in the top center in Figure 3A. When the V_rel_ value is 0, it signifies that the variant under study is unable to incorporate the modified dATP or exhibit any measurable enzymatic activity.

At position 408, eight mutations (A/I/Y/M/P/V/S/C) were identified that showed catalytic activity for modified dATP incorporation, shown in Figure 3A (left). L408I is depicted in the top left of Figure 3A in the active site structure. Only two mutations at position 409, A and G, showed enzymatic activity, with the V_rel_ of mutation of the natural Y to A (Figure 3A, center) being higher than that to G. In our test, smaller amino acids such as alanine and glycine displayed the strongest positive effects at this position, suggesting it has a potential role as a steric gatekeeper in KOD pol. Residues that function as steric gatekeepers in the DNA polymerase family are typically highly conserved, and this phenomenon is also evident in other B-family DNA polymerases. For example, in 9 °N DNA polymerase, which belongs to the family B DNA polymerases, the conserved steric gate residue is Y409^48,49^. At position 410, nine mutations (I/C/S/V/P/L/M/G/A) exhibited an increased value of V_rel_ (Figure 3A right), including the original residue of the WT enzyme, P410 (Figure 3A above right), which exhibited the best performance in incorporating modified dATP, on the background of Mut_1. Most variants at position 485 exhibited an increased V_rel_ compared to Mut_1, shown in Figure 3B. Among these, six mutants, including F/I/Y/L/K/E, displayed values of V_rel_ more than three times higher than Mut_1, with E exhibiting the highest enzymatic activity. Residues L/E/A at position 485 are shown in the active site structure in the top of Figure 3B. A previous study highlighted that A485L as an essential mutation for nucleotide analog discrimination^50^. Interestingly, we found that A485E performed satisfactorily in modified dATP incorporation, possibly due to the additional mutations included in motif A (L408A, Y409A, P410A) in our study compared to previously reported polymerase variants^45^. In addition, we observed relatively stable inter-subdomain contacts between A485E and Q332 (as illustrated in Figure 3B, top center). The contacts of those two residues may provide support for conformational changes in the finger domain, allowing for a rapid transition from a closed state to an open state during the polymerization process.

By individually evaluating saturation mutations at positions 408, 409, 410, and 485, some variants with catalytic activity were obtained. These findings suggest that mutations in specific residues play a crucial role in determining the catalytic activity of KOD pol. Further investigation is necessary to explore whether there exists any potential synergy for the incorporation efficiency of modified nucleotides between those variants that were found to have catalytic activity.

Based on the screening results of the previous round of saturation mutagenesis, mutants with increased V_rel_ values at positions 408/409/410/485, were selected for the second round of combinatorial mutant library construction, shown in Figure 4A. The screening results of thirty combined mutants from the combinatorial mutant library are displayed in Figure 4B. Eight mutants showed a V_rel_ increase of more than 8-fold compared to Mut_1 (Figure 4C), among which Mut_C2 (D141A, E143A, L408I, Y409A, A485E) exhibited the highest increase in V_rel_, which was over 16-fold compared to Mut_1. In addition, most of those combinatorial mutants exhibited a higher catalytic efficiency compared to single-point mutants.

**Figure 4.**
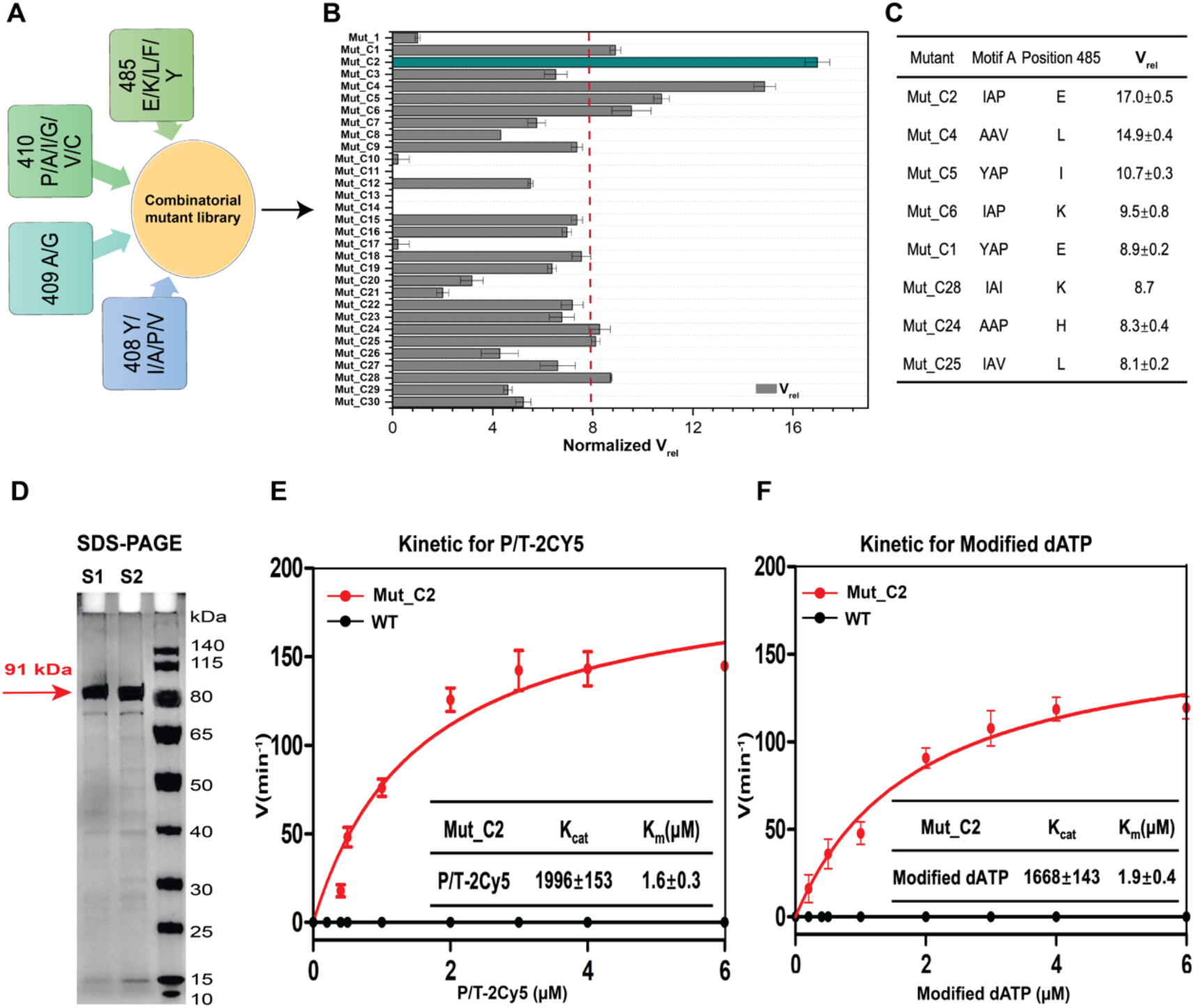
Overview of the screening results of combined mutagenesis at positions 408, 409, 410, and 485 of KOD pol. A) Schematic diagram of the combinatorial mutant library construction. The library was constructed by site-directed mutagenesis, combinatorial mutagenesis, and degenerate codon mutagenesis. Amino acid substitutions selected at position 408 include Y/I/A/P/N, at position 409 are A/G, at position 410 are P/A/I/G/V/C, and at position 485 are E/K/L/F/Y. B) displays the results of screening thirty combined mutants from the library, measuring their V_rel_ values which are shown in a bar graph. Mut_C2 (in dark green) showed the highest increase in V_rel_, over 16-fold compared to Mut_1. The other mutants are colored gray. C) The mutants resulted in a more than 8-fold increase in V_rel_ compared to Mut_1. D) SDS-PAGE analysis (12% gel) with WT KOD pol represented by S1 and KOD variant Mut_C2 represented by S2, respectively. The amount of protein loaded was around 1.0 µg, and the purification method involved lysing cells with lysozyme at 37 °C for 10 min and centrifugation after heating at 80°C for 30 min. The gel was run at 120V voltage and 1-2 h running time at room temperature. E) and F) The kinetic test results of Mut_C2 and WT KOD pol towards variations of P/T-2Cy5 (E) and modified dATP (F) under the otherwise same conditions. The experimental data were fitted non-linearly using the Michaelis-Menten equation with GraphPad Prism 5, and the values of k_cat_ and K_m_ are presented in the table insert the graph. Each kinetic experiment was performed in triplicate.

In order to further assess the catalytic efficiency of Mut_C2, tailored kinetic assays were developed for P/T-2Cy5 and 3’-O-azidomethyl-dATP labeled with a Cy3 dye (modified dATP), separately. These kinetic assays were conducted with varying concentrations of either P/T-2Cy5 or modified dATP ranging from 0 to 6 µM while maintaining the other one at a constant concentration of 2 µM. Fundamental kinetic parameters k_cat_ and K_m_ were obtained from nonlinear regression of the Michaelis-Menten equation. The kinetic results of Mut_C2 are shown in Figures 4E and 4F. The ratio of k_cat_/K_m_ for the variation of P/T-2Cy5 was 1263.3, and the ratio of k_cat_/K_m_ for the variation of modified dATP was 887.2. Notably, Mut_C2 exhibited similar performance in both types of substrate variations, indicating an improved efficiency in incorporating modified dATP into the DNA strand. The kinetic data of WT KOD pol shows only a flat line (Figures 4E and 4F), indicating that it failed to incorporate modified dATP, as expected. Mut_C2 was selected as the parent sequence for the next round of mutagenesis.

Following the evaluation of residues situated in the catalytic active center, our next step is to conduct a further assessment with a focus on the critical residues present in the DNA strand binding region of KOD pol.

### Computational screening

The identification of pivotal residues implicated in DNA binding can facilitate subsequent enhancements in DNA polymerase catalytic activity. To achieve this goal of identifying key residues involved in the DNA strand binding region of KOD pol, we selected 93 specific amino acid residues, most of them located within a 4 Å distance from the DNA strand in the crystal structure of KOD pol (PDB ID: 5MOF) (Figure 5A). These residues were chosen based on their potential to interact with the DNA through hydrogen bonds, salt bridges, and other types of interactions critical for DNA replication. The selected residues are located across critical domains of KOD pol, including thumb, finger, and palm regions, which play essential roles in DNA replication (Figure 5A). Previous studies reported that the binding process between DNA polymerase and the DNA strand with the correct dNTP pairing is slower than the chemical incorporation process^51^. Raper *et al.* proposed that the nucleotide binding and incorporation must be much faster than the binding equilibration of a polymerase and DNA (E + DNA ⇌ E·DNA)^52^, indicating the binding of DNA polymerase and DNA strands may be a rate-limiting step in the whole process. Therefore, mutations at these selected positions will affect the binding between KOD pol and DNA strands, either increasing or decreasing the binding rate, which could in turn affect the overall efficiency of KOD pol for polymerizing unnatural nucleotides. Mutagenesis in the DNA strand binding region provides valuable insights for a comprehensive understanding of the mechanism underlying the polymerization of modified nucleotides by KOD pol.

**Figure 5.**
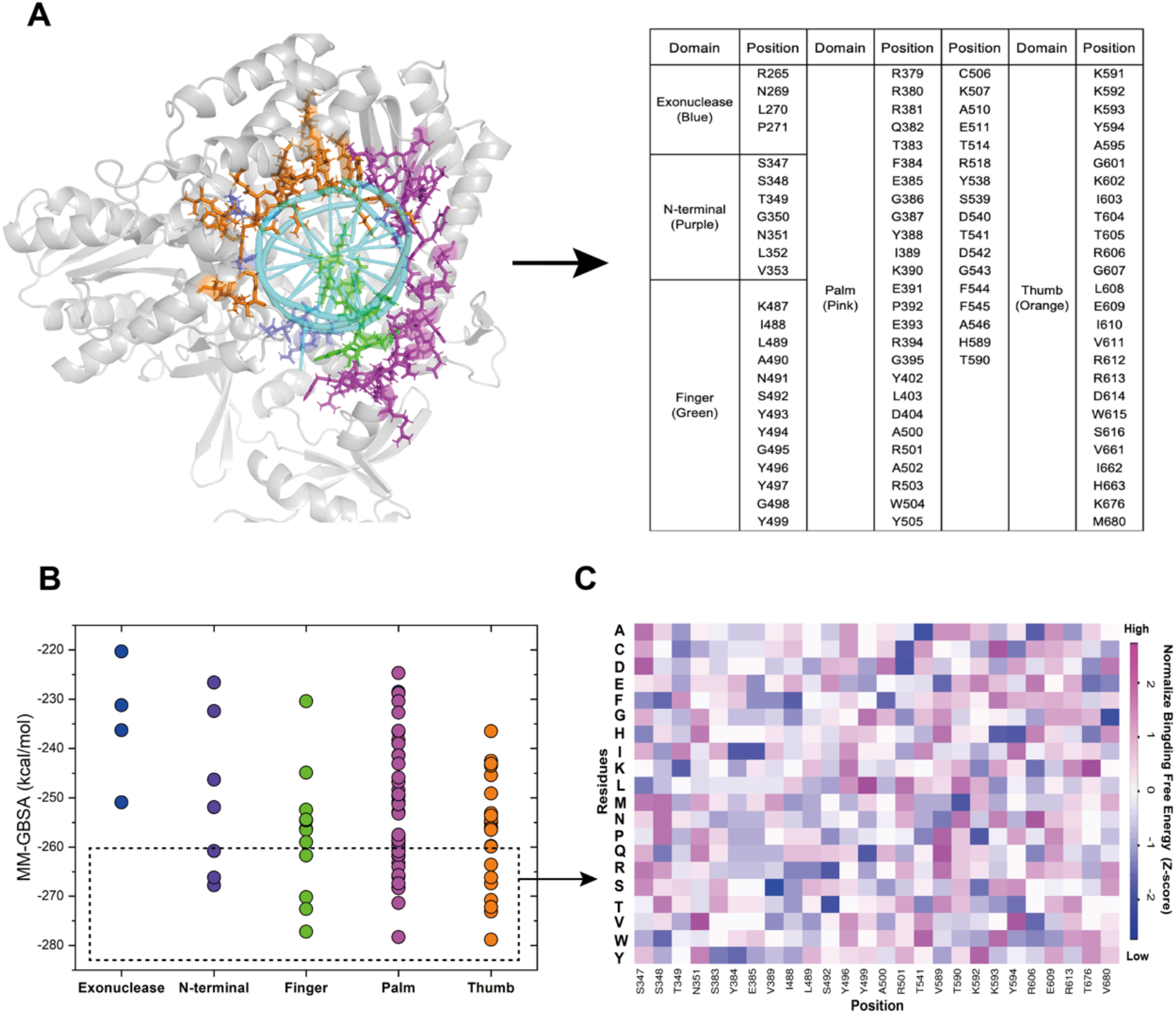
Results of computational screening of residues involved in polymerase binding to dsDNA. A) The location of 93 specific amino acid residues, most of them located within a 4 Å distance from the DNA strand in the crystal structure of KOD pol (PDB ID: 5MOF). The corresponding residues are shown as sticks and color-coded on the structure: blue for the Exo domain, purple for N-term, green for finger, pink for palm, and orange for thumb subdomain. The rest of the protein structure is shown in cartoon format and colored grey. The DNA strand is shown in cartoon format and colored cyan. The right panel displays a table presenting all residues that were selected for MD simulation. B) The average binding free energy value of the 93 residues calculated by MM-GBSA, with each region color-coded accordingly. The residues within the black dashed box were predicted to exhibit stronger binding affinity. C) Heatmap of the binding free energy of 26 distinct mutation sites selected from panel B (dashed box).

Virtual saturation mutagenesis was performed for the selected 93 residues using molecular dynamics simulation and the binding energy with dsDNA was predicted for each mutant. The workflow of simulation and calculation involving a method called MM-GBSA is presented in Supplementary Figure 2 and Supplementary Figure 3. We developed an automatic processing pipeline in Python to perform this virtual screening, including mutant pretreatment, molecular dynamics simulation, and binding energy calculation (Supplementary Figure 2). The average binding energy value of saturation mutations of the 93 residues, obtained from the MM-GBSA method, is plotted in Figure 5B and listed in Supplementary Table 1. All these residues with an average computed binding energy < -260 kcal/mol, which was lower than that of WT KOD pol, were selected for further investigation (Figure 5B, dashed box). Some interesting positions previously identified by Kropp *et al*^10^, including residues 349, 383, 389, 541, 592, and 606, were also included. The resulting next-round library consisted of 26 distinct mutation sites, and their binding energy was visualized as a heat map in Figure 5C. Variants with more negative computational free energy values are expected to exhibit stronger binding affinities to dsDNA. Finally, we selected two variants at each site with the most negative binding free energies for further experimental screening.

### Mutant screening of the third-round mutagenesis

The third-round library consisting of 52 mutants containing the 26 critical amino acids identified in virtual screening was constructed by site-direct mutagenesis. Mut_C2 (D141A, E143A, L408I, Y409A, A485E) which exhibited the best performance in the second-round screening, was used as the parent or starting variant for constructing the library in this round. The enzymatic activity of these mutants was quantified using equation 2, which is similar to equation 1 except that Mut_C2 serves as the reference. The V_rel_ of Mut_C2 was therefore 1.

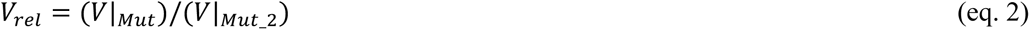

The kinetic performance of KOD variants is described by the Michaelis-Menten parameters k_cat_ and K_m._ The relative kinetic performance of those mutants is characterized by the ratio of k_cat_/K_m_|_Mut_ relative to that of Mut_C2. k_cat_/K_m_|_Mut_ and k_cat_/K_m_|_Mut_C2_ were measured and calculated in the same way, with varying P/T-2Cy5 ranging from 0 to 6 µM while maintaining the modified dATP at a constant concentration of 2 µM. We define the relative kinetic performance of the mutant variants as E(Mut), as shown in Equation 3. The subscript after the vertical bar “|” indicates the mutant used for measurement. The Mut_C2 variant served as the reference, with a catalytic efficiency (E(Mut)) value of 1.

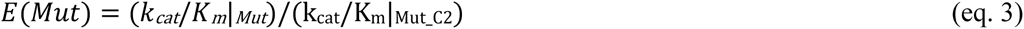

The enzyme activity of 52 variants was screened under the same experimental conditions as described above, and their V_rel_ values were calculated and compared (Figure 6 A). 14 variants displayed a V_rel_ value greater than that of Mut_C2, with one variant, Mut_D34 (V589H), exhibiting an over 2-fold greater V_rel_ to Mut_C2. We selected 39 mutant variants, with a V_rel_ value higher than 0.5, to perform further kinetic screening measurements. Although some mutant variants did not show significantly higher enzyme activity than Mut_C2, we still selected them because they contain mutations located close to the DNA strand in the structure of KOD pol.

**Figure 6.**
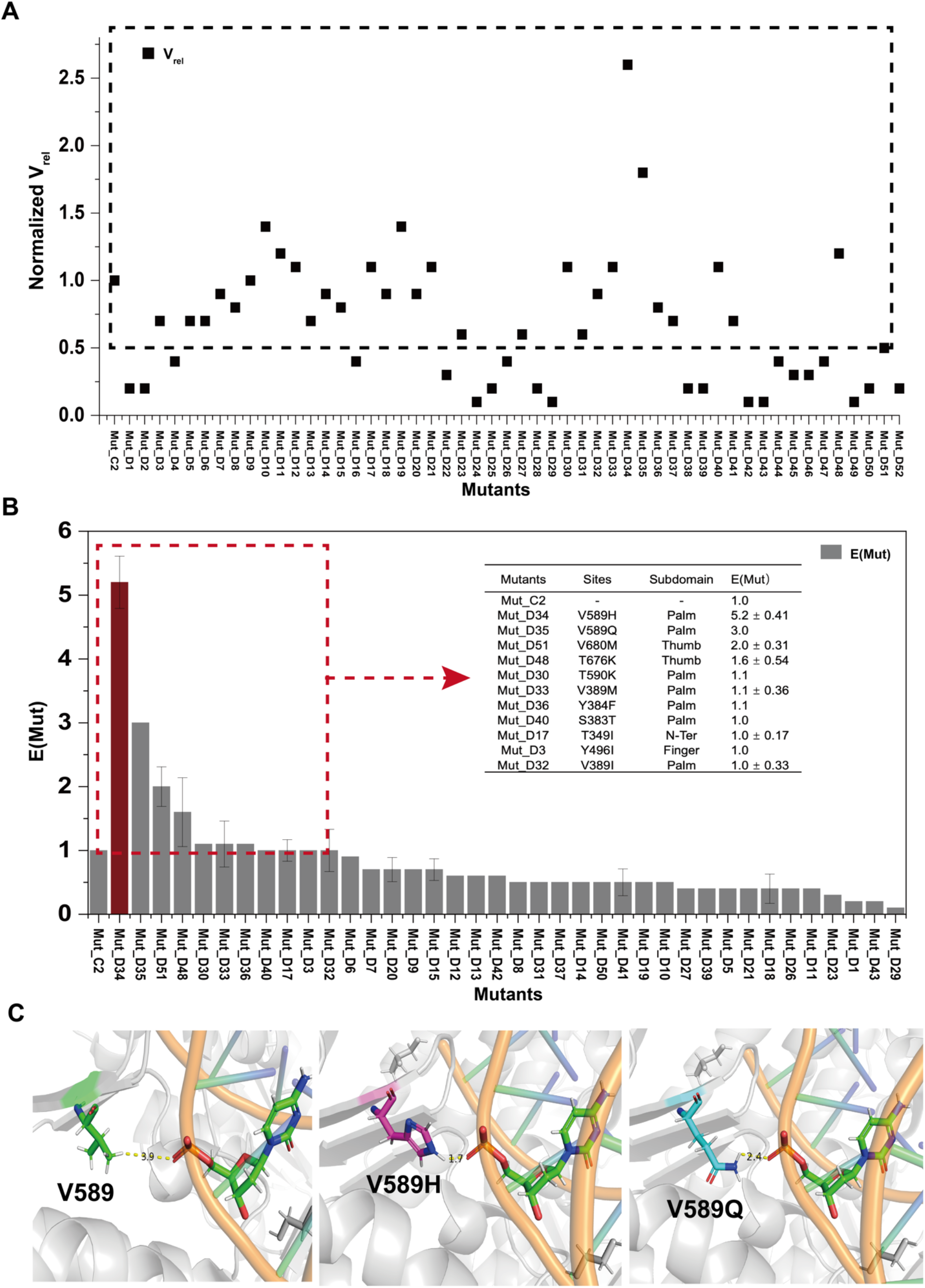
Screening results of third-round variants constructed based on Mut_C2. A) The enzyme activity screening results of these variants. The V_rel_ (black square) of these variants was calculated according to equation 2. The variants with a V_rel_ > 0.5 are included inside the dashed box. B) The relative kinetic screening results of those variants with a V_rel_ >0.5 are shown in the dashed box in Panel A. The kinetic parameters of the Michaelis-Menten equation, such as k_cat_ and K_m_, were calculated by analyzing the data using GraphPad Prism 5 software. E(Mut) of these mutants was calculated according to equation 3. Mut_D34 corresponds to the red column, while the other variants are shown as grey bars. B) The site information, location, and kinetic results of variants with E(Mut) > 1. C) The location of residue 589 with V/H/Q placed in the crystal structure of KOD pol (PDB ID: 5MOF). The DNA strand is depicted in a cartoon format, with the backbone-colored orange, while one of the deoxyribonucleotides is shown as sticks and colored by element. The residues at position 589 are displayed as sticks and colored green for V (in the left panel), magenta for H (in the middle panel), and cyan for Q (in the right panel). Atom contacts identified between residues (V, H, Q) at position 589 and the DNA strand are illustrated as an example. The values of V_rel_ and E(Mut) were summarized in Supplementary Table 2. The distances predicted by the PyMOL software are represented by dashed yellow lines.

E(Mut) values of these variants were determined and presented in Figure 6 B. Among this set, 11 variants displayed more favorable E(Mut) values than Mut_C2, listed in the table embedded in Figure 6 B. Mut_D34, which contains the V589H mutation, outperformed the other variants with the highest observed E(Mut) value of 5.2. Mut_D35, carrying the V589Q mutation with no charged side chain, showed improved kinetic activity (E(Mut) = 3.0) compared to Mut_C2 (V589). The observed increase in catalytic activity in Mut_D34 and Mut_D35 can be attributed to the presence of V589H and V589Q mutations, which are located in the palm domain of KOD pol. The distances of atom contacts between residues V, H, and Q at position 589 and the DNA strand in the crystal structure of KOD pol (PDB ID: 5MOF) are shown in Figure 6C, which are 3.9 Å, 1.7 Å, and 2.4 Å, respectively. A closer distance implies a higher possibility of hydrogen bonds forming, which can subsequently impact the binding between DNA and the protein. In addition, we also speculated that the positively charged side chain of histidine (H) at position 589 increased the binding affinity between KOD pol and the DNA strand, facilitating the incorporation process by virtue of strong and favorable electrostatic interaction with the double-stranded DNA. This has been supported by previous studies^39^. For example, Kropp *et al.* reported that the structure of KOD pol has a long crack with an electro-positive potential extending from the DNA binding region at the thumb domain upwards along the β-hairpin and the palm domain to the N-terminal domain^11^. The increased positive charge introduced in the DNA binding region of the polymerase results in enhanced affinity towards the template DNA, leading to a more stable binding^53^. Mut_D51, which carried the V680M mutation, and Mut_D48 carrying the T676K mutation, demonstrated 2-fold and 1.6-fold improvements in E(Mut) compared to Mut_C2, respectively. Among the investigated mutation sites, five were located in the palm domain (V589H/Q, T590K, V389I, S383T, Y384F), two in the thumb domain (T676K, V680M), one in the finger domain (Y496I), and one in the N-terminal region (T349I). Among the mutation sites investigated in this study, residues 383/384/389 were located in the loop of the KOD pol structure, while residues 589/590/676/680 contributed to the binding between KOD pol and the DNA strand. The screening results showed that those mutated residues either directly or indirectly influence the binding of KOD pol and the DNA strand, leading to an increase in the incorporation efficiency during polymerization. Therefore, we performed further combinations with these well-performing mutants to explore possible interactions and synergies between the identified sites. To accomplish this, we generated stepwise combination variants based on Mut_D34 exhibited the best performance in this round of screening.

### Stepwise combination of effective mutations

Based on the last-round screening results, the best-performing variant Mut_D34 was selected as the parent for a fourth round of combinational mutagenesis to further improve the catalytic activity. To comprehensively assess the catalytic performance of stepwise combined variants, we conducted kinetic assays towards both substrates of P/T-2Cy5 and modified dATP. These assays allowed us to evaluate the kinetic performance of each new variant individually with respect to the DNA strand and modified dATP. The relative kinetic performance was evaluated using Equation 4, which is similar to Equation 3. In this round of screening, the Mut_D34 variant served as a reference, with a catalytic efficiency (E(Mut)) value of 1.

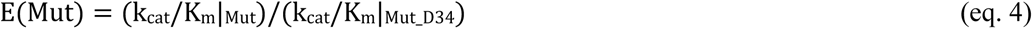

To distinguish the relative kinetic performance of two substrates, we used E(Mut)/P/T-2Cy5 to represent the kinetic measurement towards varying P/T-2Cy5. Similarly, E(Mut)/modified dATP represents the kinetic measurement towards varying modified dATP.

The screening results for E(Mut)/P/T-2Cy5 of combined variants are shown in Figures 7A and 7B. We combined V680M (Mut_D51) with the mutations of variant Mut_D34, generating the combination variant Mut_E1 which displayed slight improvement in E(Mut), confirming that the V680M mutation stacked on Mut_D34 improved catalytic performance. We then added T676K (Mut_D48) to Mut_E1, which introduced a positive charge located on the thumb domain. The results demonstrated that these three mutation sites displayed positive interactions for improving polymerase catalytic efficiency. The resulting improved variant was designated as Mut_E2. Further single site mutations were introduced to Mut_E2, including T590K, V389H, Y496L, T349F, E385M, and Y384F. Four mutants carrying specific residues, namely Mut_E3 (Y349F), Mut_E4 (Y384F), Mut_E5 (Y496L), and Mut_E8(V389I), displayed increased E(Mut) compared to Mut_E2. We then generated a two-point and three-point superposition of S383T, Y384F, and V389I, leading to the variants Mut_E9 and Mut_E10 with further improved kinetic performance. The E(Mut) value of Mut_E9, which carries the mutations S383T and Y384F, showed a 3-fold increase compared to Mut_E1 and approximately a 1.6-fold increase compared to Mut_E2. The E(Mut) value of Mut_E10 carrying S383T, Y384F, and V389I, showed a 4.5-fold increase compared to Mut_E1. However, attempts to further combine Mut_E10 with Y349F and Y496L did not result in an improved variant.

**Figure 7.**
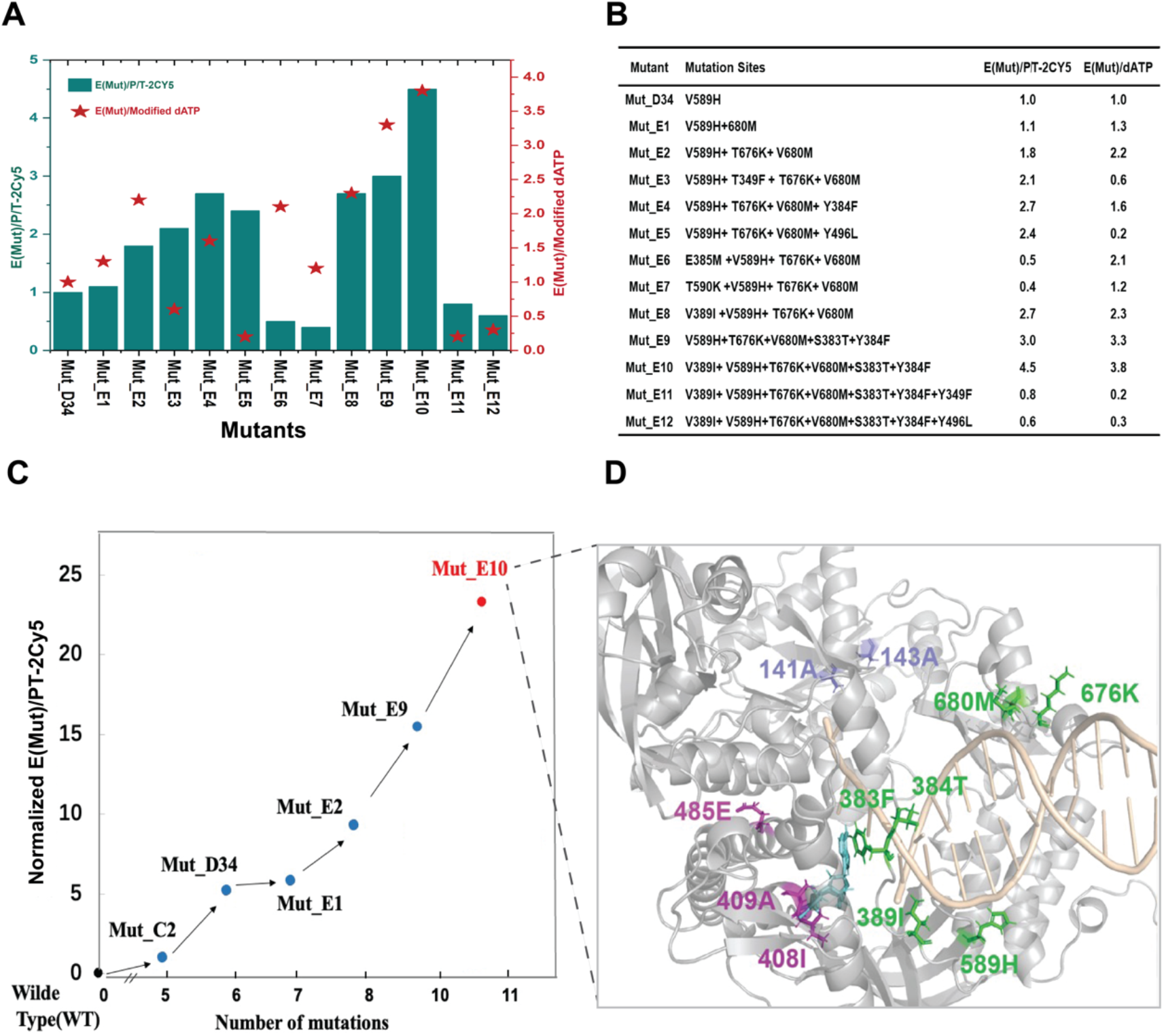
Screening results of stepwise combinational mutagenesis. Mut_D34 was used as the parental variant in this round, and additional mutation sites were introduced using site-directed mutagenesis. A) Relative kinetic screening results for variants, which include their kinetic performance towards P/T-2CY5 (E(Mut)/P/T-2Cy5, displayed as green bars), as well as their performance towards modified dATP (E(Mut)/modified dATP, displayed as red stars). The kinetic assays were conducted for 1-2 hours at 40°C with varying concentrations of P/T-2CY3 or modified dATP, using excitation at 530 nm and detection at 676 nm. B) The mutation sites information and the corresponding dynamic results of combined variants. C) Illustration of the evolution process of Mut_E10 (red spots), which comprises 11 mutation sites compared to the wild-type KOD pol. During its semi rational evolution process, key mutants (black spots) include Mut_C2, Mut_D34, Mut_E1, Mut_E2, and Mut_E9. D) The locations of eleven mutation sites of the variant Mut_E10 are shown in the crystal structure of KOD pol (PDB ID: 5MOF). These residues are shown as sticks, the rest of the protein structure is shown in cartoon format and colored grey. The sites located in the exonuclease region, including D141A and E143A, are colored light blue. The sites associated with the catalytic activity center, including L408I, Y409A, and A485E, are colored light magenta. The sites related to DNA binding, including S383T, Y384F, V389I, V589H, T676K, and V680M, are colored green. The DNA strand is shown in cartoon format and colored in light wheat. The 3’-O-azidomethyl-dATP is shown in sticks and colored cyan.

E(Mut)/modified dATP values of these mutants were calculated and are also shown in Figures 7A and 7B. Interestingly, we found that some variants behaved inconsistently in kinetic experiments against P/T-2Cy5 and modified dATP. For example, Mut_E3 and Mut_E5 exhibited a two-fold increase compared to Mut_D34 in the P/T-2Cy5 kinetic assay but displayed poorer kinetic performance in the modified dATP. Except for some variants that exhibit inconsistent performance with P/T-2Cy5 and modified dATP, most exhibit consistent behavior. Notably, Mut_E10 exhibited the highest performance for variations of both substrates, indicating a high level of efficiency in incorporating modified dATP into the DNA strand. Overall, our data show that introducing combinatorial mutations at specific positions within the DNA-binding region of KOD pol significantly affects its efficiency in incorporating modified nucleotides during the polymerization process.

The whole semi-rational evolution process leading to the identification of Mut_E10, is depicted in Figures 7C and 7D. With two-round libraries screening, we obtained a five-mutation variant Mut_C2 (D141A, E143A, L408I, Y409A, A485E). Mut_C2 demonstrated a higher catalytic efficiency in incorporating modified dATP compared to wild-type KOD pol. We then performed virtual screening and selected several mutation sites located in the DNA binding region for further investigation. By combining the V589H mutation with the mutations present in Mut_C2, we obtained a variant called Mut_D34. Mut_D34 exhibited a 5.2-fold improvement in E(Mut)/PT-2Cy5 compared to Mut_C2. Based on Mut_D34, we conducted stepwise combinatorial mutation screening and ultimately obtained a best-performance variant Mut_E10. Mut_E10 exhibited an exceptional catalytic efficiency, with a relative increase of over 23-fold in E(Mut)/PT-2Cy5 compared to Mut_C2. This indicates a significant improvement in the polymerase’s ability to incorporate modified dATP. Our results highlight the importance of combinatorial mutations of multiple amino acids in improving the incorporation efficiency of KOD pol for modified nucleotides.

### Sequencing performance validation of KOD variants in NGS

In order to examine whether the observed improvement in the polymerization of Mut_E10 brought significant values in kinetic performance in sequencing applications, we tested the performance of Mut_E10 in different NGS platforms. Since the launch of the Human Genome Project, next-generation sequencing (NGS) platforms have had a very significant impact on biological research^54^. We first compared the sequencing performance of Mut_E10 to that of WT and Mut_C2 on the BGISEQ-500 platform with SE50 sequencing, which can assess the relative improvements achieved through the use of the engineered variants. Furthermore, we conducted a further sequencing evaluation of Mut_E10 on the MGISEQ-2000 platform, which involved PE100 sequencing. We performed rigorous purification for WT, Mut_C2, and Mut_E10. The protein purity of each sample was confirmed to exceed 95% through SDS-PAGE analysis, shown in Supplementary Figure 4.

The sequencing results of WT KOD pol, Mut_C2, and Mut_E10 on the BGISEQ-500 platform are presented in Figure 8. The BGISEQ-500 platform featuring cPAS and DNA Nanoballs (DNB™) technology is a widely used NGS platform^55^. The library used for sequencing was *E. coli*, the number of sequencing cycles was SE50, and the temperature was 55 °C. We compared the performance of the three different enzymes, WT KOD pol, Mut_C2, and Mut_E10, by replacing the original enzyme in the supplied sequencing kit using the BGISEQ-500 sequencing platform. Notably, the WT KOD pol group did not show any bright spots on the chip (Figure 8A), indicating that no fluorescently labeled probes were extended. In contrast, the Mut_C2 group presented bright spots, indicating the successful extension of modified probes, with the bright spots representing the incorporated fluorescently labeled probes in the DNB event, as shown in Figure 8A. Then, the sequencing results of Mut_E10 and Mut_C2 were analyzed and compared. The quality distribution of the sequencing reads, including Q30, ESR, and mapping rate, represents the overall sequencing quality of the enzyme-replaced platforms, shown in Figures 8B and 8D. The results revealed that Mut_E10 exhibited superior performance compared to Mut_C2 which has an apparent decrease after 20 cycles. A higher Q value indicates a lower probability (P) of base misidentification^56^. Mut_E10 had a higher Q30 (> 93%) throughout the entire cycle, making it highly suitable for NGS applications. The data revealed that Mut_E10 exhibited superior performance in terms of lag % and runon % compared to Mut_C2 (Figure 8C). These parameters reflect the incorporating ability of Mut_E10 as the sequencing enzyme in the NGS application. Based on these results, we conclude that Mut_E10 is a promising candidate for the BGISEQ-500 sequencing platform. Subsequently, we investigated the performance of Mut_E10 on another sequencing platform, CoolMPS^TM^ sequencing.

**Figure 8.**
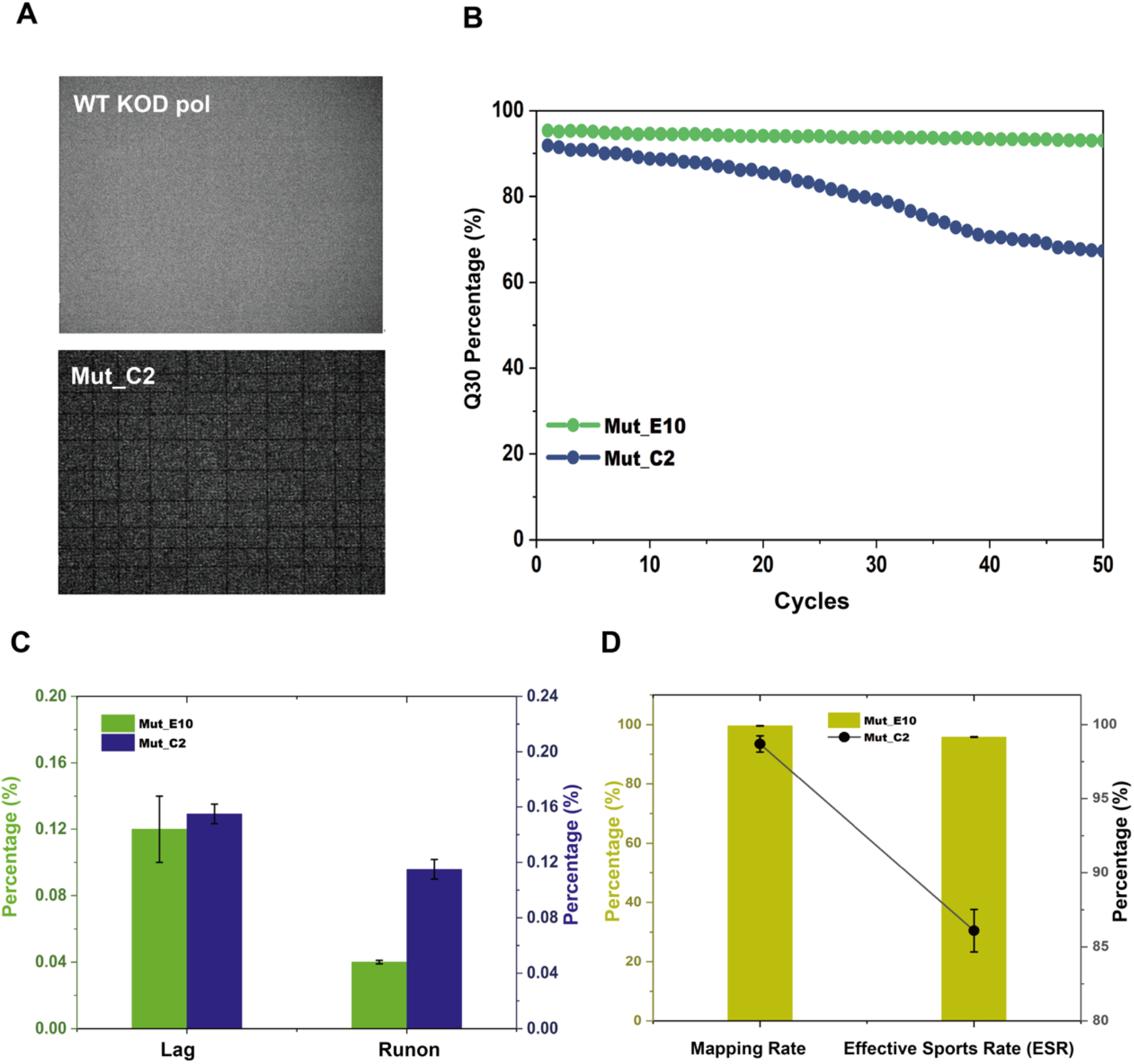
Sequencing results of WT KOD pol, Mut_C2, and Mut_E10 on the BGISEQ-500 platform. A) Representative images in the sequencing process. During the sequencing process, DNA polymerases polymerize modified nucleotides, and then remove excess fluorescence before imaging the chip. The WT KOD pol group showed no bright spots on the chip, indicating no fluorescently labeled probes were extended. In contrast, the Mut_C2 group exhibited bright spots, indicating the successful extension of modified probes. The area of a small square in the chip shown in the diagram is 0.882*0.882 mm. B) Quality portion distribution on reads (Q30) for Mut_C2 (blue line) and Mut_E10 (green line). Mut_E10 had a stable curve and performed better in each sequencing cycle in comparison to Mut_C2. C) Lag and Runon comparison for Mut_C2 (blue) and Mut_E10 (green) in sequencing results. Mut_E10 had lower values, indicating better extension efficiency. D) Mapping rate and ESR comparison between Mut_C2 (black squares) and Mut_E10 (olive). Mut_E10 had a higher mapping rate and ESR, suggesting better sequencing quality.

Furthermore, we applied Mut_E10 on the MGISEQ-2000 platform featuring CoolMPS^TM^ technology for PE100 sequencing. CoolMPS^TM^ is a novel massively parallel sequencing chemistry, which was developed by Drmanac’s group at MGI^57^. The sequencing experiments were performed in duplicates on a single chip in Lane1 and Lane2. The library used for sequencing was *E. coli*, the temperature was 55 °C, and the sequencing process was carried out following the published protocols^58^. The significant sequencing results were listed in a concise table in Figure 9A. Both the first and second strands exhibited high mapping rates exceeding 99%. The ESR value was reported to be above 78%, and the average error rate was less than 0.16, further indicating the accuracy and quality of the sequencing data. During the base-calling of sequencing, the Q30 score remained consistently higher than 95% for the first-strand sequencing and a gradual decrease in Q30 was observed for the second-strand sequencing (Figure 9B), which is considered acceptable for PE100 testing. The average lag % and runon % values for both strands were comparatively low and are depicted in Figures 9C and 9D. In addition, Korostin reported that Illumina HiSeq 2500 and MGISEQ-2000 have similar performance characteristics after comparing both whole-genome sequencings (PE150) in terms of sequencing quality, number of errors, and performance^58^. Our sequencing data also exhibited comparable performance to that of HiSeq 2500, including high mapping rates (>99 %) and Q30 values (>97 %), with Q30 being around 96%. The sequencing data obtained is satisfactory for the MGISEQ-2000 sequencing platform, according to the reported range of NGS analysis generated^52,56^. Taken together, these results demonstrate that Mut_E10 is suitable for practical applications in the MGISEQ-2000 platform.

**Figure 9.**
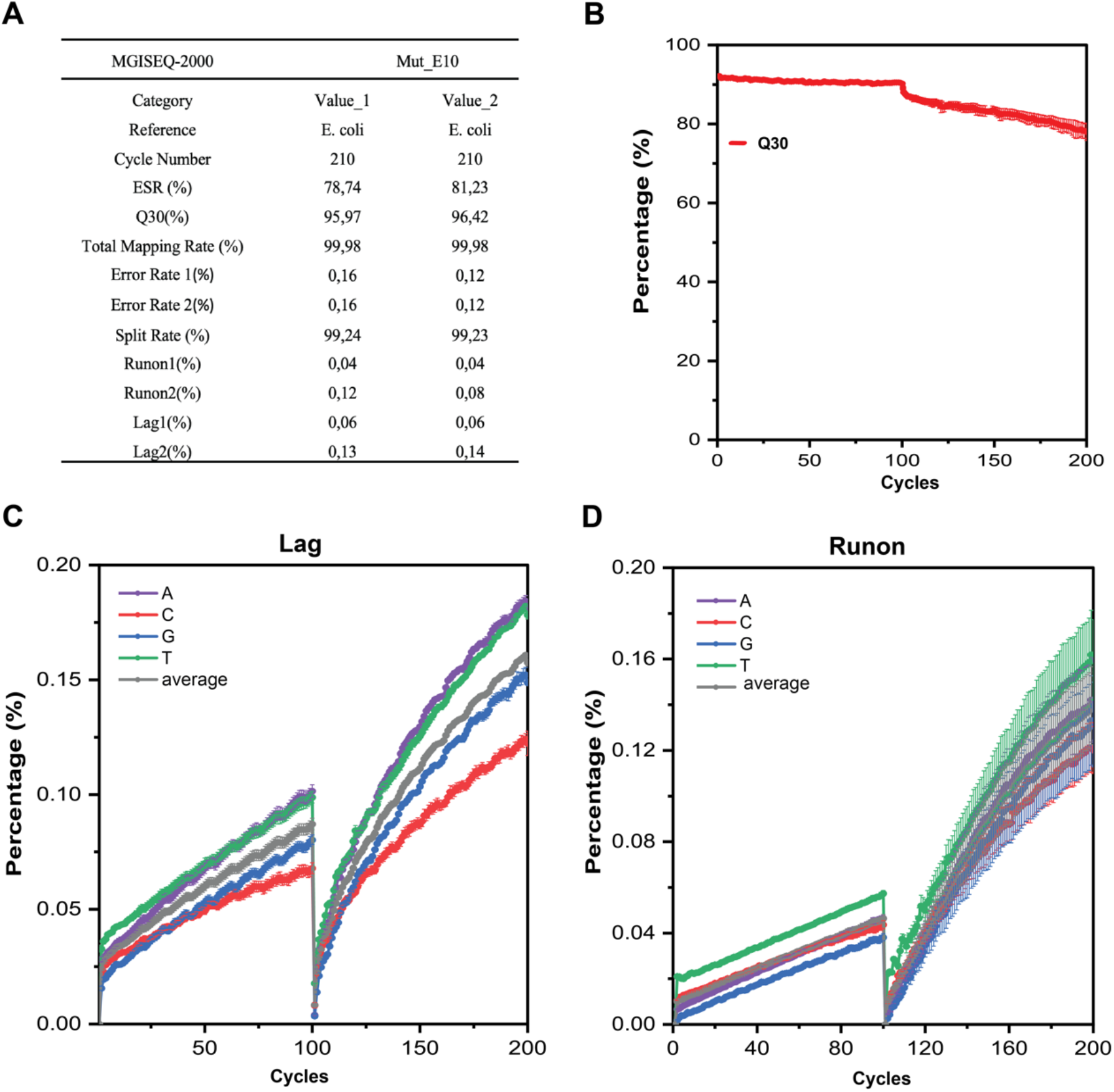
Sequencing results of Mut_E10 on the MGISEQ-2000 platform. A) Overview of the significant sequencing information and major sequencing quality metrics or data, with PE100 sequencing. Note: Error Rate: base error rate; Split Rate: barcode split rate. Value_1 and Value_2 are duplicates of sequencing data obtained from Lane 1 and Lane 2, respectively, on the same chip during sequencing. B) The average Q30 value of Value_1 and Value_2 during sequencing in panel A), which serves as an indicator of base calling accuracy. C) and D) show the average lag and runon percentages of Value_1 and Value_2 during sequencing in panel A), respectively. These values serve as indicators of the completeness of the polymerization reaction that occurred within the DNB.

Finally, considering that the enzyme in the original sequencing reagent has been directly replaced, it is believed that there is still potential for enhancing the sequencing quality of Mut_E10 on both sequencing platforms. One potential avenue for improvement lies in optimizing the sequencing buffer to further enhance the performance and accuracy of the sequencing process^59^. Consequently, the sequencing quality achieved with Mut_E10 on both the BGISEQ-500 and MGISEQ-2000 platforms fell within the acceptable range comparable to the “gold standard” set by NGS analysis generated using the Illumina platform^58^.

## Discussion

Mut_E10 displayed satisfactory performance on two types of sequencing platforms including BGISEQ-500 and MGISEQ-2000, as indicated by our sequencing data. These results suggest that Mut_E10 possesses the potential for commercialization and is adaptable to different sequencing platforms. To our knowledge, the cPAS technology of the BGISEQ-500 platform utilized 3’-O-blocked reversible terminators labeled with four different fluorescent dyes (four colors)^9,60^. On the other hand, the CoolMPS^TM^ technology of the MGISEQ-2000 platform achieved sequencing using monophosphate nucleotides with a 3’-O-azidomethyl blocking group and an NHS linker on phosphate^61,62^, which can rapidly bind (within 30 seconds) with nucleobase-specific fluorescently labeled antibodies^57^. Previous studies reported DNA polymerases can incorporate dNMPs at apurinic/apyrimidinic (AP) sites or similar damaged sites but with generally low efficiency and a propensity for error insertion^63,64^. Remarkably, Mut_E10 exhibits high efficiency in incorporating both chemically modified dNTPs and dNMPs. The incorporation efficiency of unnatural nucleotides depends on the linker used to anchor the modification to the nucleobase, especially with sterically demanding groups^33^. Therefore, we speculate that the steric size requirements for the incorporation of chemically modified dNTPs and dNMPs by DNA polymerases may be similar in this study. However, the mechanisms responsible for the incorporation of both unnatural substrates in these two sequencing platforms, are yet to be fully understood. Our analysis focuses on the potential causes or mechanisms by which these identified mutations improve the polymerization efficiency of Mut_E10. Therefore, we conducted further analysis on these identified mutations. Except for the two sites (D141A, E143A) in the exonuclease region, the remaining nine mutation sites can be categorized into three distinct categories.

The first category of mutational sites includes those that affect the binding of Mut_E10 polymerase with the DNA strand. These sites are S383T, Y384F, V389I, V589H, T676K, and V680M, as shown in Figure 7D. MM-GBSA calculations indicated that modifications at these sites enhanced the ability of Mut_E10 to bind the DNA strand, accelerating DNA strand capture. Notably, residues 589H and 676K possess positive charges and are expected to exhibit higher binding affinities for negatively charged DNA when the polymerase binds to the DNA strand. Residues 383T, 384F, and 389I are located near the DNA strand and nucleotides that form the back of the active pocket likely contribute positively to the formation of a stable active pocket. Furthermore, residues 383T, 384F, 389I, and 676K are situated on the coil structure. To avoid disrupting the original coil structure, we screened for mutants that were similar to the original residues. Interestingly, experimental results revealed that if the properties of the residue side chains at these points changed significantly, the calculated binding energy might be stronger, but the experimental test results did not provide any indications for an increased binding affinity (data not shown). These findings indicate that a stable coil or loop structure of the DNA polymerase plays an important role in modified nucleotide polymerization activity. Excessive and disruptive changes to these residues on the coil structure are thus not recommended.

The second category of sites includes those where mutations altered the dNTP catalytic pockets, such as L408I and Y409A, as shown in Figure 7D. The modified dATP structure (with an azido methyl terminal group at the 3’ end, a long linker, and Cy3 fluorescent group) is depicted in Figure 1B. When the modified dATP with a blocking group enters the catalytic pocket, its blocking group would collide with the surrounding L408,Y409,P410 residue side chains, as shown in Figure 7D. The mutations in motif A (L408I/Y409A/P410) of Mut_E10 have smaller side chains, thus provide more space to accommodate the incoming modified dATP with blocking group, making Mut_E10 more active for modified dATP incorporation. Interestingly, Mut_1 with motif A (L408A/Y409A/P410A) should have even more space in this area for modified dATP, however, the catalytic activity of this variant was not optimal. There may be other critical factors that should be considered affecting the catalytic efficiency of DNA polymerase besides the space in motif A, such as metal ions. For example, metal ions can affect the incorporation efficiency of DNA polymerase for dNMPs or dNTPs^65^. Further study could consider the interaction between metal ions and crucial residues. Overall, our findings suggest that when engineering DNA polymerase to incorporate a modified substrate with a larger group, using small side chain residues in the catalytic pockets is recommended.

In the third site category, A485E mutation affects the conformational change of the finger domain of KOD pol, as shown in Figure 7D. The binary complex structure of KOD pol that bound only to the DNA strand was compared with the ternary complex structure of KOD pol that bound to DNA complex and dATP, as depicted in Figure 2A. Previous reports have suggested that the movement of the finger domain may be advantageous for the fidelity of KOD DNA polymerase^66^. In Mut_E10, the A485 residue on the rotating alpha helix of the finger domain was substituted with A485E, and this resulted in the formation of a salt bridge (as illustrated in Figure 3B, top center) with Q332 on the opposite helix of the exonuclease domain, thereby preventing the rotation of the finger domain. Evidently, this mutation had a detrimental effect on the catalytic activity of KOD pol when natural nucleotides were used. However, in the case of modified nucleotides employed in NGS, which possess long linkers and fluorescent groups, the locking of the finger domain has the potential to enhance the stability of the modified substrate access channel. This improvement can lead to an increased entry and exit rate of modified nucleotides from the catalytic pocket. The experimental outcomes indicated that this mutation significantly improved the enzyme’s catalytic efficiency.

In addition, the semi-rational evolution strategy employed for enhancing the efficiency of modified nucleotide incorporation by KOD pol exhibited satisfactory efficiency compared to some reported methods^66,67^. Kennedy *et al*. conducted a steady-state kinetic analysis to examine the DNA synthesis process by DNA polymerase using natural or modified nucleotides^67^. They employed denaturing polyacrylamide gel electrophoresis to distinguish and assess the extension of individually modified nucleotides in an exogenous primer. In contrast, our approach utilizes fluorescence as an indicator for kinetic assays, offering advantages such as higher throughput, increased sensitivity, and a more streamlined analysis process. Nikoomanzar *et al*. developed a microfluidic-based deep mutational scanning method to evolve a replicative DNA polymerase (KOD pol) from *Thermococcus kodakarensis* for TNA synthesis^66^. The authors identified positive sites in the finger subdomain of KOD pol through a single round of sorting from 912 mutants and subsequently combined them step-by-step to obtain a double mutant variant. In contrast, our screening strategies achieved higher screening efficiency, resulting in Mut_E10 containing nine mutation sites (excluding 141A/143A), with fewer experimentally validated mutants (not exceeding 150). Nevertheless, our screening method using a microplate reader does not possess a comparable level of throughput as their approach, which employed a microfluidic-based droplet screening strategy.

Therefore, the screening method for future studies could consider employing microfluidic-based droplet screening combined with FRET fluorescence as an indicator, as this approach has the potential to overcome the limitations in screening throughput. For instance, we tested a small fraction of the variants from the computational library and combinatory library, and future studies could employ higher-throughput screening methods to assess more variants that may have been missed during the evolutionary process. Furthermore, further exploration of residues for continually improving catalytic efficiency of KOD pol can be conducted around the amino acid residues neighboring these key positions, such as motif B including residues 484/485/486, and residues around 389/589/676/680.

## Conclusion

To conclude, this study presents a comprehensive workflow for engineering WT KOD DNA polymerase to incorporate unnatural nucleotides suitable for NGS. The workflow can be summarized into five stages: construction of the screening strategy, identification of residues in the active pocket for improvement, identification of residues located in the DNA strand binding region for improvement, step-wised combination, and characterization of variants in a specific application. The variant Mut_E10, which had eleven mutation sites, was found to enable successful compatibility between the two sequencing platforms, namely BGISEQ-500 and BGISEQ-2000. These beneficial mutation sites identified in this study could act as inspiration for the engineering and improvement of other archaeal B-family DNA polymerases. In addition, the successful engineering of KOD DNA polymerase with improved catalytic efficiency for unnatural nucleotides proves the great potential of the enzyme engineering strategy constructed, which offers a new solution of polymerase engineering to fulfill versatile applications.

## Methods and Materials

### Reagents and instruments

The primer and template labeled at the 5’-terminus by Cy5 dye were ordered from Invitrogen. The primer-template (P/T-2Cy5) was prepared by mixing the primer and template at a 1:1 molar ratio and annealing at 80 °C for 10 min and then cooling down to room temperature in a water bath. The sequence information of P/T - 2Cy5 is shown in Figure 1C. The primers designed with mutations were ordered from Genscript Biotech. 3’-O-azidomethyl-dATP-Cy3 (modified dATP) was supplied by the BGI synthetic chemistry group, and the structure was shown in Figure 1B. Potassium phosphate, MgSO_4_, KCl, (NH4)_2_SO_4_, Tris, NaCl, HCl, EDTA, NaOH, imidazole, glycerol, lysozyme, LB broth, kanamycin, PMSF, and IPTG were purchased from Thermo Fisher. 12% SDS-PAGE gel and gel running buffer were purchased from Bio-Rad. Affinity chromatography columns (HisTrap HP 5ml column) and ion exchange columns (HiTrap Q FF 5mL and HiTrap SP HP 5ml column) were purchased from GE Healthcare. The ÄKTA Pure Protein Purification system was purchased from GE Healthcare. The microplate reader was purchased from Bio Tek.

### Mutation library construction

The codon-optimized gene encoding this exonuclease-deficient KOD pol (named Mut_1) was synthesized by Genscript Biotech and then cloned into vector pD441 with kanamycin resistance. The corresponding expressed protein features a 6xHis tag at the N-terminus. The first mutant library consisted of single-site saturation mutagenesis at four positions: 408, 409, 410, and 485. The saturation mutagenesis at positions 408, 409, 410, and 485 was performed based on the background of triple alanine substitutions at the other three positions. The site-directed mutagenesis kits were purchased from Thermo Fisher. The PCR reaction conditions were as follows: one cycle at 98 °C for 10 s; 20 cycles at 98 °C for 10 s,72 °C for 2.5 min, followed by elongation at 72 °C for 5 min and hold at 4 °C. The PCR products were visualized by 1.2 % agarose gel electrophoresis and ultraviolet light. Then, 1 μL of DpnI (NEB) was added to the PCR mix and incubated for 2 h at 37 ℃. 5μL of the digested PCR product was transformed into the 50 μL *E. coli* DH5α competent cells (Tiangen). Recombinant plasmids were prepared by using the QIAprep Spin Miniprep Kit (QIAGEN) and confirmed by sequencing analysis. 1 μL of each correct plasmid was transformed into 50 μL *E. coli* BL21(DE3) competent cell (Tiangen) for protein expression.

### Protein expression and purification

1 mL of LB medium in a 96-deep-well plate was used for small-scale expression, and 1 L of LB medium in a conical flask was used for large-scale expression. Cells grown in LB medium containing 50 μg/mL kanamycin were induced at OD600 nm = 0.6 - 0.8 by the addition of 0.5 mM IPTG. Protein production was carried out at 25 °C, 220 rpm overnight.

Semi-purification was performed using cell pellets collected from 1 mL culture in a 96-deep-well plate. The cell pellets were resuspended in Buffer 1 (20 mM Tris-HCl, 10 mM KCl, 10 mM (NH4)_2_SO_4_, 0.1% Triton, 4 mM MgSO_4_, 1.25 mM PMSF, and 1 mg/mL lysozyme, pH 7.6) with a ratio of 0.04 g cell pellets per mL buffer. The resuspended cells were incubated at 37 °C for 10 minutes, followed by thermal denaturation at 80 °C for 30 minutes. The supernatant containing proteins was collected after centrifugation at 12,000 X g for 20 minutes. The protein concentration was measured and estimated by using the Bradford protein assay^68^.

We rigorously purified mutant proteins for use on NGS platforms and the purification methods as the following process. For protein purification, cell pellets collected from 1 L cultivation were resuspended in Buffer 2 (500 mM NaCl, 5 % glycerol, 20 mM imidazole, 1.25 mM PMSF, 50 mM potassium phosphate, pH 7.4) and then disrupted with a high-pressure homogenizer AH-1500 purchased from ATS engineering company. After a 30-minute thermal denaturation at 80 °C, the centrifugation at 12,000 rpm 4 ℃ (Beckman Avanti J-26) for 30 mins was performed to remove cell debris. After centrifugation, the supernatant was filtered through a 0.22 µm membrane filter. The subsequent purification process was performed using ÄKTA pure 25 (GE Healthcare) with Nickel affinity chromatography (5 mL HisTrap HP column, GE Healthcare), anion ion exchange chromatography (5 mL HiTrap Q FF column, GE Healthcare), and cation ion exchange chromatography (5 mL HiTrap SP HP column, GE Healthcare), sequentially. For the affinity purification process, the supernatant was loaded onto the pre-equilibrated column with a flow rate of 2.5 mL/min so that the retention time will be 2 min. A wash step using 50 mL Buffer 2 with a flow rate of 5mL/min was added before elution. The elution procedure was performed with a linear gradient of Buffer 3 (500 mM NaCl, 5% glycerol, 500 mM imidazole, 50 mM potassium phosphate, pH 7.4) from 0 to 100% in 10 CV with a flow rate of 5 mL/min. Fractions with 280 nm UV signal above 100 mAu were collected. The collected samples from Ni purification were diluted 6-fold with Buffer 4 (5% glycerol, 25 mM potassium phosphate, pH 6.6) to decrease the concentration of NaCl to 500 mM. The diluted sample was loaded onto the pre-equilibrated HiTrap Q FF 5 mL column (GE Healthcare) at a flow rate of 2.5 mL/min. The flowthrough sample was collected and loaded onto the pre-equilibrated HiTrap SP HP 5 mL column (GE Healthcare) with a flow rate of 2.5 mL/min. The HiTrap SP HP 5 mL column was pre-equilibrated with Buffer 5 (50 mM NaCl, 5 % Glycerol, 50 mM Potassium Phosphate, pH 7.4). The elution procedure was performed with a linear gradient of Buffer 6 (1 M NaCl, 5 % Glycerol, 50 mM Potassium Phosphate, pH 6.6) from 0 to 60% in 10 CV. The first peak eluted was collected. Finally, the collected sample was dialyzed with Buffer 7 (20 mM Tris, 200 mM KCl, 0.2 mM EDTA, 5 % glycerol, pH 7.4) at 4 °C overnight and then stored at -20 °C with 50 % glycerol. Protein concentration was determined by measuring the absorbance at 280 nm using a microplate reader (Bio Tek) and calculated using the extinction coefficient of 1.393 M^-1^ cm^-1^ predicted by the ExPASy server. The purity of the protein was analyzed using 12% SDS-PAGE.

### Enzyme activity screening of KOD variants

The polymerization reaction mixture contained 0.1 μM P/T-2Cy5, 0.25 μM modified dATP, 2.5 μM native dC/G/TTP each, 2 μg BSA, 1X KOD Screening Buffer (20 mM Tris-HCl, 10 mM KCl, 10 mM (NH_4_)_2_SO_4_, 0.1% Triton, 4 mM MgSO_4_, pH 8.8) and 0.5 μg KOD protein. The reaction was initiated by adding each semi-purified KOD mutant protein. The enzyme activity assay was performed in 384-well (black clear-bottom) plates (Corning®) at 40 °C for 1-2 hours. The FRET Cy5 fluorescence signal was measured as relative fluorescence units (RFUs) in appropriate time intervals by exciting at 530 nm and detecting the emission at 676 nm. Measurements were performed with a gain setting at 60 and shaking for 10 seconds before measurement in a microplate reader (Bio Tek). The enzyme activity (reaction rate V) was defined as the maximum slope of the FRET Cy5 signal, expressed as RFUs/min. The data were analyzed and plotted using OriginPro.

### Enzyme kinetic assays

The reaction mixture contained 10 μM native dC/G/TTP each, 2 μg BSA, 1X KOD Screening Buffer (20mM Tris-HCl, 10 mM KCl, 10 mM (NH_4_)_2_SO_4_, 0.1 % Triton, 4 mM MgSO_4_, pH 8.8) and 0.5 μg KOD protein. The kinetic assays were carried out with varying concentrations of P/T-2Cy5 or modified dATP ranging from 0 to 6 µM, while the concentration of modified dATP or P/T-2Cy5 remained constant at 2 µM. The reaction was initiated by adding KOD mutant protein. The assay was also performed in 384-well (black clear-bottom) plates (Corning®) at 40 °C for 1-2 hours. The FRET Cy5 fluorescence signal was measured by exciting at 530 nm and detecting the emission at 676 nm. Measurements were performed with a gain setting at 60 and shaking for 10 seconds before measurement in a microplate reader (Bio Tek). The maximum slope (reaction rate V) of the FRET Cy5 signal for each concentration of P/T-2Cy5 was calculated and used for determining the kinetic parameters (k_cat_ and K_m_). The kinetic data were fitted non-linearly using the Michaelis-Menten equation with GraphPad Prism 5 to obtain k_cat_ and K_m_.

### MM-GBSA screening method

The complex structure of KOD pol and dsDNA adopts the 5OMF PDB structure. Additional small molecules in both structures were removed, and the systems were energy minimized in 80 mM sodium chloride aqueous solution, followed by NVT ensemble MD relaxation of 50 ns^69^. We used the Modeler 9.19 software to carry out saturation mutagenesis at 93 residues on the polymerase and then processed all 1860 mutants uniformly. Due to this large number of calculations, we designed an automatic processing workflow in Python. In an 80 mM sodium chloride aqueous solution, five rounds of energy minimization were first carried out for the structure generated by homologous modeling. In the five rounds of simulations, the binding coefficient to heavy atoms of protein and DNA was gradually weakened from 80 kcal / (mol · Å) to 0 kcal / (mol · Å). After that, NVT ensemble MD was carried out at 350 K, and finally, NPT ensemble MD of 10-25 ns was carried out at 310 K. We sampled the trajectory of the system that reached equilibrium in the last part and calculated the binding energy by the MM-GBSA method^70^.

### Sequencing in the NGS platform

BGISEQ-500 is a desktop platform developed for DNA or RNA sequencing by BGI. It utilizes DNA NanoBalls (DNBs) technology (DNBSEQ^TM^). The dNTPs used in DNBSEQ^TM^ technology were modified with four different fluorescent dyes in the base and a reversible blocking group in its 3’-side. To evaluate the sequencing performance of Mut_E10 in the BGISEQ-500 platform, an *E. coli* library prepared by following the instructions described in the manufacturer’s use manual of MGIEasy PCR-Free DNA Library Prep Kit (PN: 1000013452, MGI-tech) was used to make DNB following BGISEQ-500 protocol^60^. The DNBs were then loaded into the patterned nanoarrays. The SE50 sequencing kit was supplied by the BGI sequencing group. WT KOD pol, Mut_C2, and Mut_E10 were used to replace the original enzyme in the sequencing kit and performed sequencing tests in the BGISEQ-500 sequence platform employing the SE50 mode. Base calling and mapping were performed as previously described^55^.

MGISEQ-2000 is a high-throughput platform developed by MGI. The sequencing performance of Mut_E10 was also tested in the DNBSEQ-2000 platform employing CoolMPS^TM^ technology. The cold nucleotides used in CoolMPS^TM^ technology are monophosphate nucleotides with a 3’-O-azidomethyl blocking group. CoolMPS^TM^ PE100 High-throughput Sequencing kit (PN: 1000016935) was purchased from MGI-tech. The same E. coli library used for BGISEQ-500 was the same for MGISEQ-2000. The sequencing process followed the instructions described in the MGISEQ-2000 High-throughput Sequencing Set User Manual^57^. Mut_E10 was used to replace the original enzyme in the CoolMPS^TM^ PE100 sequencing kit. Mut_E10 was used to sequence the flow cells loaded with DNBs prepared with the E. coli library in different runs. Base calling and mapping steps were performed as reported previously^58^.

### Sequence information

>Seq_1(WT KOD DNA pol)

MILDTDYITEDGKPVIRIFKKENGEFKIEYDRTFEPYFYALLKDDSAIEEVKKITAERHGTVVTVKRVEKVQKKFLGRPVEVWKLYFTHPQDVPAIRDKIREHPAVIDIYEYDIPFAKRYLIDKGLVPMEGDEELKMLAFAIATLYHEGEEFAEGPILMISYADEEGARVITWKNVDLPYVDVVSTEREMIKRFLRVVKEKDPDVLITYNGDNFDFAYLKKRCEKLGINFALGRDGSEPKIQRMGDRFAVEVKGRIHFDLYPVIRRTINLPTYTLEAVYEAVFGQPKEKVYAEEITTAWETGENLERVARYSMEDAKVTYELGKEFLPMEAQLSRLIGQSLWDVSRSSTGNLVEWFLLRKAYERNELAPNKPDEKELARRRQSYEGGYVKEPERGLWENIVYLDFRSLYPSIIITHNVSPDTLNREGCKEYDVAPQVGHRFCKDFPGFIPSLLGDLLEERQKIKKKMKATIDPIERKLLDYRQRAIKILANSYYGYYGYARARWYCKECAESVTAWGREYITMTIKEIEEKYGFKVIYSDTDGFFATIPGADAETVKKKAMEFLKYINAKLPGALELEYEGFYKRGFFVTKKKYAVIDEEGKITTRGLEIVRRDWSEIAKETQARVLEALLKDGDVEKAVRIVKEVTEKLSKYEVPPEKLVIHEQITRDLKDYKATGPHVAVAKRLAARGVKIRPGTVISYIVLKGSGRIGDRAIPFDEFDPTKHKYDAEYYIENQVLPAVERILRAFGYRKEDLRYQKTRQVGLSAWLKPKGT*

## Acknowledgment

This study was supported by Shenzhen Engineering Laboratory Molecular Enzymology (Fa Gaishen [2018] No. 958). W.L. Thanks to the National Natural Science Foundation of China, China Grant No. 21505134. Alexander K. Buell thanks the Novo Nordisk Foundation for support (NNFSA170028392). Thanks to China National GeneBank DataBase & BGI’s Sequencing Platform and MGI’s CoolMPS^TM^ platform.

## Author Contributions

Conceived and designed the experiments: Lili Zhai, Yue Zheng. Performed the experiments: Lili Zhai. Performed the simulation: Zi Wang. Performed the application test: Liu Fen, Jingjing Wang, Hongyan Han. Analyzed the data: Lili Zhai. Contributed reagents/materials/analysis tools: Qingqing Xie, Yue Zheng, Wenwei Zhang, Yuliang Dong. Wrote the paper and SI: Lili Zhai, Zi Wang. Reviewed and revised the paper: Lili Zhai, Yue Zheng, Qingqing Xie, Chongjun Xu, and Alexander Kai Buell.

