## Supplemental information for "Semi-rational evolution of a recombinant DNA polymerase for modified nucleotide incorporation efficiency"

### Supplementary Information

#### Experimental validation

We conducted a validation of our screening method, using the identified mutants as an exemplar. In order to validate our screening method, we utilized wild-type KOD pol and two mutants (Mut_1 and Mut_C2) with different enzymatic activity from the first-round screening as examples selected from the initial screening due to their distinct catalytic efficiencies toward the modified dATP. In this approach, we performed a quantitative assessment of the enzyme activity of the KOD variants, which was crucial for evaluating their catalytic performance.

After semi-purification, mutant proteins were analyzed by SDS-PAGE gel electrophoresis, and the estimated protein purity was approximately 80 % based on ImageJ analysis of the gels, as shown in Supplementary Figure 1A. In order to quantify the mutant protein, we established a standard curve using bovine serum albumin (BSA) as a standard and measured the absorbance at 595 nm through the Bradford assay[38,39]. The BSA concentration range was between 0 to 2.0 µg/µL, as shown in Supplementary Figure 1B. Through linear regression analysis of the standard curve, we were able to consistently quantify the KOD mutant protein. After calculating the total protein concentration (Conc. Total) using the Bradford method, the final target protein concentration was determined by multiplying the Conc. Total by 80 %. This approach enabled us to determine the protein concentration of KOD mutants obtained in the previous step and ensure their purity for subsequent analysis and characterization.

We calculate the maximum slope of the increased fluorescence signal (FRET Cy5) as a surrogate for the enzyme activity of KOD mutants. To assess the FRET Cy5-based enzymatic activity of the KOD mutants, we added 0.5 µg of each mutant protein to the reaction system, with the reaction buffer serving as the negative control (no enzyme added). As shown in Supplementary Figure 1C, a steady starting was observed before the rise of the FRET Cy5 fluorescence signal, which could be attributed to a temperature increase during the reaction. WT KOD DNA polymerase failed to generate a FRET Cy5 fluorescence signal, indicating WT

KOD pol cannot catalyze modified dATP into the DNA strand. In contrast, Mut_C2 exhibited a higher catalytic efficiency compared to Mut_1, generating more FRET Cy5 fluorescence signals within the same time frame. This observation suggests that Mut_C2 is capable of efficiently incorporating modified dATP into the DNA strand, making it a promising candidate for further characterization and development. Overall, our experimental screening approach was effective in distinguishing different levels of enzyme activities of the mutants.

In addition, fluorescently labeled substrates used in our experiments were at risk of fluorescence quenching during storage and usage, which raises concerns about the reliability and reproducibility of data obtained from experiments utilizing fluorescent labeling, particularly in situations where extended storage or multiple freeze-thaw cycles are involved. To get rid of the influence of fluorescence quenching, we preferred using the direct readout of the fluorescent signal values in each batch test, not the absolute substrate concentrations. Positive control was employed in each experiment as a reference for all the other tested mutants. By using this compromise solution, the efficiencies of mutants tested not in the parallel experiments can be compared. During every round of the evolution process, the parent mutant was utilized as a positive control.

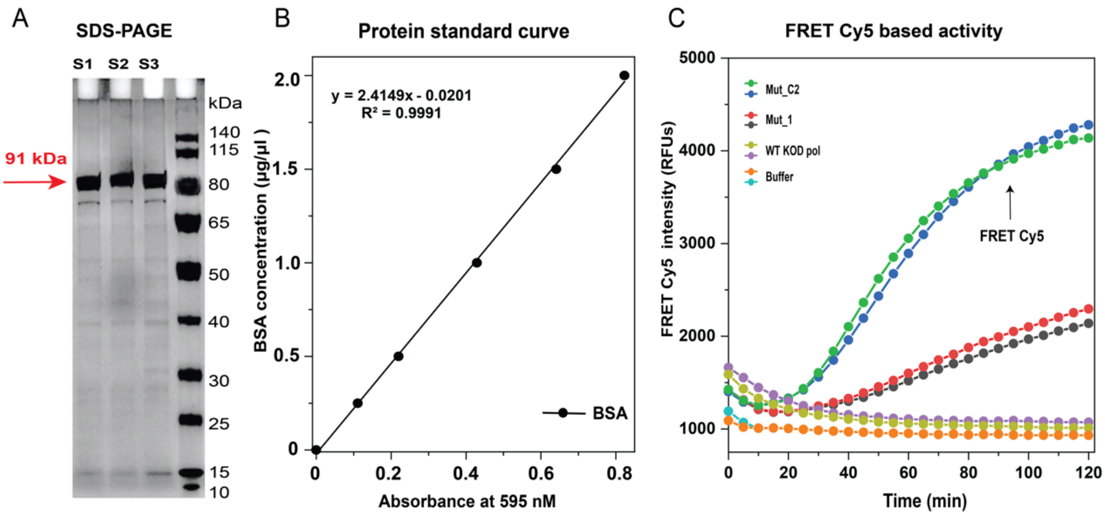


Supplementary Figure 1. Experimental validation of the reliability of the screening methods. A) This figure displays the SDS-PAGE analysis (12% gel) with WT KOD pol represented by S2, KOD variant Mut_1 represented by S1, and KOD variant Mut_C2 represented by S3. The volume of samples loaded was around 1 μg, and the purification method involved lysing cells with lysozyme at 37 °C for 10 min and centrifugation after heating at 80°C for 30 min. The gel was run at 120V voltage and 1-2 h running time at room temperature. B) The protein standard curve was generated using bovine serum albumin (BSA) as the protein standard. Absorbance was measured at 595nm using a microplate reader (BioTek)

at room temperature after the mixture 15 minutes of incubation in the dark. C) FRET Cy5-based enzyme activities of wild-type KOD pol (olive and purple) and KOD variants Mut_1 (red and black), Mut_C2 (green and blue), and buffer (orange and cyan) are presented. In these experiments, 0.5 µg protein was used for measurements. The reactions were performed at 40°C for 2 hours with excitation at 530 nm and detection at 676 nm using a microplate reader (BioTek). The data were directly output using the Gene5 software of the microplate reader. The curve was plotted using OriginPro. Each experiment was performed by duplicates under the same conditions.

### Computational simulation

We then calculated the changes in binding free energy between the mutants and dsDNA using the MM-GBSA method. The automation process of the MM-GBSA calculation and MD simulation is shown in Supplementary Figure 2 and Supplementary Figure 3. The average binding energy of 93 residues was calculated and is shown in Supplementary Table 1. Due to the high standard deviation of MM-GBSA calculation, during data analysis, we first calculate the average value of the calculation results of 20 mutants at a single site and then select some mutants for further experimental testing to determine their lower average value. We expressed and tested several enzyme mutants with an average binding energy of < 260 kcal/mol and some interesting positions. These successfully constructed and expressed mutants were subjected to measurement of enzymatic activity and polymerization kinetics towards the DNA primer-template, and the results are shown in Supplementary Table 2.

Supplementary Table 1. Average Binding Free Energy (Kcal/mol) of 93 Mutation Sites in DNA-binding region.

| Mutant | Avg. (Kcal/mol) | Mutant | Avg. (Kcal/mol) | Mutant | Avg. (Kcal/mol) | Mutant | Avg. (Kcal/mol) |
|---|---|---|---|---|---|---|---|
| R265 | -236.3 | P392 | -228.8 | W504 | -264 | G601 | -256.3 |
| N269 | -231.2 | E393 | -259.5 | Y505 | -260 | K602 | -255.1 |
| L270 | -250.9 | R394 | -258.9 | C506 | -249.6 | I603 | -254.6 |
| P271 | -220.3 | G395 | -263.2 | K507 | -230.4 | T604 | -242.5 |
| S347 | -272.9 | Y402 | -260.1 | A510 | -251.3 | T605 | -270.8 |
| S348 | -252.3 | L403 | -241.4 | E511 | -243.1 | R606 | -242.0 |
| T349 | -235.1 | D404 | -258.8 | T514 | -232.7 | G607 | -259.8 |
| G350 | -260.8 | K487 | -256.3 | R518 | -246.8 | L608 | -256.2 |
| N351 | -271.6 | I488 | -231.9 | Y538 | -261.6 | E609 | -273.1 |
| L352 | -251.9 | L489 | -244.9 | S539 | -266.1 | I610 | -272.2 |
| V353 | -246.3 | A490 | -277.2 | D540 | -260.9 | V611 | -243.5 |

| | | | | | | | |
|---|---|---|---|---|---|---|---|
| R379 | -278.3 | N491 | -270.2 | T541 | -257.5 | R612 | -253.6 |
| R380 | -252.8 | S492 | -239.6 | D542 | -268.1 | R613 | -267.4 |
| R381 | -236.6 | Y493 | -256.4 | G543 | -258.6 | D614 | -263.6 |
| Q382 | -238.3 | Y494 | -256.5 | F544 | -257.5 | W615 | -259.9 |
| T383 | -255.7 | G495 | -254.6 | F545 | -268.4 | S616 | -278.8 |
| F384 | -228.5 | Y496 | -261.7 | A546 | -245.9 | V661 | -243.1 |
| E385 | -228.5 | Y497 | -252.4 | H589 | -265.7 | I662 | -249.1 |
| G386 | -271.4 | G498 | -259.0 | T590 | -267.4 | H663 | -256.2 |
| G387 | -241.3 | Y499 | -254.4 | K591 | -243.2 | K676 | -256.2 |
| Y388 | -260.7 | A500 | -250.2 | K592 | -245.4 | M680 | -253.1 |
| I389 | -238.3 | R501 | -249.0 | K593 | -255.3 | | |
| K390 | -250.8 | A502 | -248.4 | Y594 | -256.0 | | |
| E391 | -258 | R503 | -253.2 | A595 | -253.1 | | |

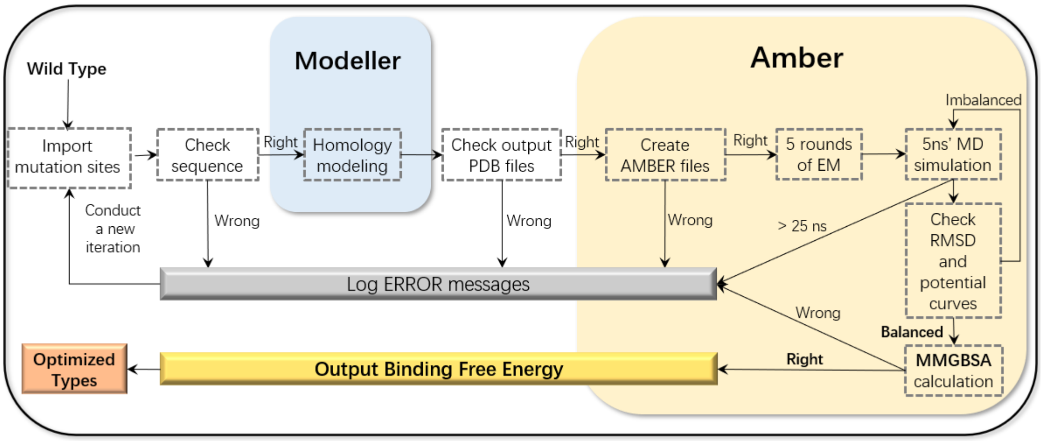


Supplementary Figure 2. Automation process of MM-GBSA calculation. EM signifies Energy Minimization; MD signifies Molecular Dynamics; RMSD signifies the Root-Mean-Square Deviation.

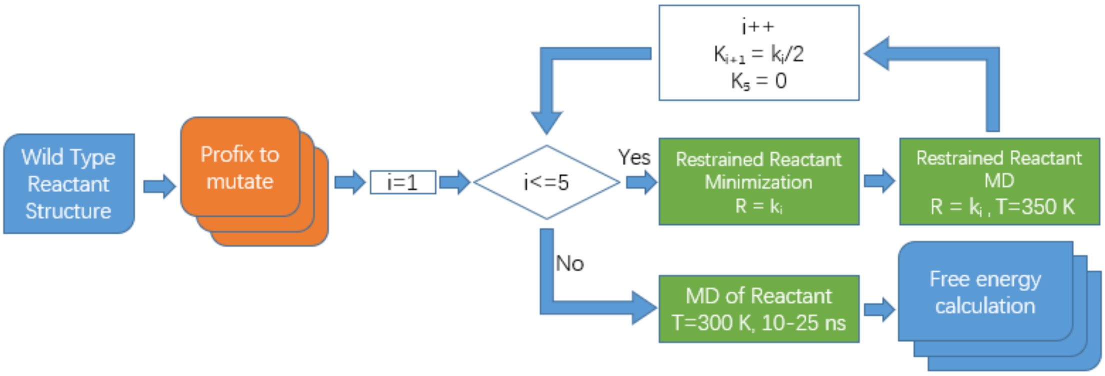


Supplementary Figure 3. Mutants' pretreatment and MD simulation process. i was the cycle number of energy minimization; $k_i$ was the restraint coefficient; T was temperature.

### Gel analysis

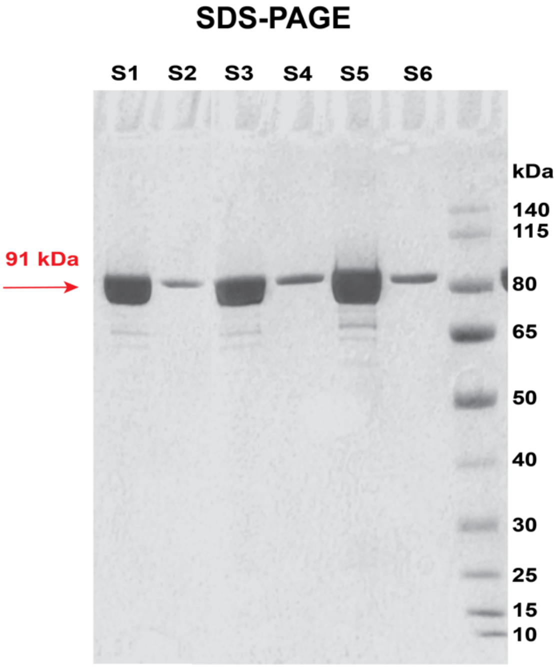


Supplementary Figure 4. The SDS-PAGE analysis (12% gel) for rigorously purified proteins with WT KOD pol represented by S1 (5 μg) and S2 (0.25 μg), KOD variant Mut_C2 represented by S2(5 μg) and S4 (0.25 μg), and KOD variant Mut_E10 represented by S5 (5 μg) and S6 (0.25 μg). We employed a three-step purification process involving Ni affinity chromatography and anion-cation exchange chromatography using ÄKTA. The estimated protein purity was approximately 95 % based on ImageJ analysis of the gels.

### Data

Supplementary Table 2. Kinetic data towards the P/T-2Cy5 of combined mutants. All the mutants in this table also share the same mutation sites as Mut_C2.

| Mutants | Addition Sites | Subdomain | $V_{rel}$ [(1)] | E(Mut)/P/T-2Cy5 [(2)] |
|---|---|---|---|---|
| Mut_C2 | - | - | 1.0 | 1.0 |
| Mut_D34 | V589H | Palm | 2.6 ± 0.21 | 5.2 ± 0.41 |
| Mut_D35 | V589Q | Palm | 1.8 | 3.0 |
| Mut_D51 | V680M | Thumb | 0.5 ± 0.07 | 2.0 ± 0.31 |
| Mut_D48 | T676K | Thumb | 1.2 ± 0.07 | 1.6 ± 0.54 |
| Mut_D30 | T590K | Palm | 1.1 ± 0.35 | 1.1 |
| Mut_D33 | V389M | Palm | 1.1 ± 0.21 | 1.1 ± 0.36 |
| Mut_D36 | Y384F | Palm | 0.8± 0.14 | 1.1 |
| Mut_D40 | S383T | Palm | 1.1 | 1.0 |
| Mut_D17 | T349I | N-Ter | 1.1 ± 0.07 | 1.0 ± 0.17 |
| Mut_D3 | Y496I | Finger | 0.7 ± 0.21 | 1.0 |
| Mut_D32 | V389I | Palm | 0.9 ± 0.07 | 1.0 ± 0.33 |
| Mut_D6 | I488Q | Finger | 0.7 ± 0.11 | 0.9 |
| Mut_D7 | L489Y | Finger | 0.9 ± 0.14 | 0.7 |
| Mut_D20 | E385M | Palm | 0.9 ± 0.14 | 0.7 ± 0.19 |
| Mut_D9 | I488L | Finger | 1 | 0.7 |
| Mut_D15 | S348T | N-Ter | 0.8± 0.14 | 0.7 ± 0.17 |
| Mut_D12 | S347I | N-Ter | 1.1 ± 0.21 | 0.6 |
| Mut_D13 | S347M | N-Ter | 0.7 ± 0.21 | 0.6 |
| Mut_D42 | E609Y | Thumb | 0.1 | 0.6 ± |
| Mut_D8 | L489K | Finger | 0.8 | 0.5 ± |
| Mut_D31 | T590L | Palm | 0.6 ± 0.14 | 0.5 |
| Mut_D37 | Y384W | Palm | 0.7± 0.14 | 0.5 |
| Mut_D14 | S348M | N-Ter | 0.9 ± 0.21 | 0.5 |
| Mut_D50 | V680D | Thumb | 0.2 | 0.5 |
| Mut_D41 | S383I | Palm | 0.7 | 0.5 ± 0.21 |
| Mut_D19 | A500G | Palm | 1.4 ± 0.14 | 0.5 |
| Mut_D10 | N351H | N-Ter | 1.4 ± 0.21 | 0.55 ± 0.04 |
| Mut_D27 | R501M | Palm | 0.6 | 0.4 |
| Mut_D39 | Y594V | Palm | 0.2 | 0.4 |
| Mut_D5 | Y499L | Finger | 0.7 ± 0.07 | 0.4 |

| | | | | |
|---|---|---|---|---|
| Mut_D21 | E385W | Palm | 1.1 ± 0.14 | 0.4 |
| Mut_D18 | A500D | Palm | 0.9 ± 0.07 | 0.4 ± 0.23 |
| Mut_D26 | R501L | Palm | 0.4 | 0.4 |
| Mut_D11 | N351Y | N-Ter | 1.2 ± 0.07 | 0.4 |
| Mut_D23 | K592Y | Palm | 0.6 | 0.3 |
| Mut_D1 | S492E | Finger | 0.2 | 0.2 |
| Mut_D43 | E609G | Thumb | 0.1 | 0.2 |
| Mut_D29 | T541W | Palm | 0.1 | 0.1 |
| Mut_D38 | Y594I | Palm | 0.2 | 0 |
| Mut_D2 | S492Y | Finger | 0.2 | NA |
| Mut_D4 | Y496L | Finger | 0.4 | NA |
| Mut_D16 | T349F | N-Ter | 0.4 | NA |
| Mut_D22 | K592L | Palm | 0.3 | NA |
| Mut_D24 | K593F | Palm | 0.1 | NA |
| Mut_D25 | K593W | Palm | 0.2 | NA |
| Mut_D28 | T541I | Palm | 0.2 | NA |
| Mut_D44 | R606H | Thumb | 0.4 | NA |
| Mut_D45 | R606N | Thumb | 0.3 | NA |
| Mut_D46 | R613G | Thumb | 0.3 | NA |
| Mut_D47 | R613T | Thumb | 0.4 | NA |
| Mut_D49 | T676Y | Thumb | 0.1 | NA |
| Mut_D52 | Y499G | Finger | 0.2 | NA |

(1): The relative enzyme activity of each KOD variant was calculated with equation 2. Enzyme activity was measured by enzyme activity screening method (details in **Materials and Methods** section).

(2): The relative kinetic performance of each KOD variant was calculated with equation 3. The kinetic was measured by enzyme kinetic assay (details in **Materials and Methods** section). NA: Measurements were not performed.

The data presented represent the mean and standard deviation values obtained from duplicated independent experiments.
